# Spatiotemporal coordination of specialized cerebellar networks supports episodic memory in older adults

**DOI:** 10.64898/2026.08.03.742395

**Authors:** Tengfei Han, Ziyun Li, Chi Zhang, Mingxi Dang, Wenhao Bai, Zaizhu Han, Yaojing Chen, Zhanjun Zhang

## Abstract

Episodic memory decline in ageing is typically attributed to the cortico-hippocampal system, yet the cerebellum’s specific contribution via cerebellar-cerebral circuits remains underexplored. We analysed task-based fMRI data from 821 older adults, including a longitudinal subset of 78 participants followed for four years. Using sparse dictionary learning on task-responsive activity, we identified three non-motor networks: a bilateral lobule VI/Crus I mnemonic-monitoring network, a right lobule Crus I/II default-mode-aligned network, and a right lobule VIIb/VIIIa evidence-evaluation network. While network recruitment varied with task demands and associated with memory performance, phase-specific static cerebellar–cerebral functional connectivity showed stable organization but limited behavioural specificity. In contrast, dynamic functional connectivity patterns effectively distinguished memory-performance groups. Crucially, greater instability in dynamic functional connectivity during encoding predicted longitudinal retrieval slowing. These findings suggest that preserved episodic memory in older adults is supported by the spatiotemporal coordination of specialized cerebellar networks with distributed cerebral networks.

## Introduction

Episodic memory supports everyday remembering and is among the cognitive functions most vulnerable to ageing^1–3^. In healthy ageing, episodic memory typically begins to decline from mid-to-late adulthood, with measurable changes often emerging between 50 and 60 years of age^4,5^. Current models emphasize that episodic encoding and retrieval depend on coordinated interactions between the hippocampus and distributed cortical networks, including medial temporal, posterior medial, parietal and prefrontal regions^6,7^. Although these frameworks have substantially advanced our understanding, they leave open whether episodic memory in older adults also depends on systems beyond the canonical hippocampal–cortical architecture.

The cerebellum is a plausible candidate for such a contribution^8^. Anatomical and physiological studies show that the cerebellum is embedded in closed-loop polysynaptic circuits with association cortices, including prefrontal, parietal, temporal regions^9–11^. Human neuroimaging further demonstrates that the cerebellar cortex contains functionally differentiated territories spanning sensorimotor, attentional, executive and transmodal domains^12–15^. These findings reframe the cerebellum from a structure primarily associated with motor control to a learning system involved in prediction, temporal sequencing, error monitoring and the coordination of distributed neural processing^16–19^.

Evidence for cerebellar involvement in episodic memory has accumulated across neuroimaging and causal-intervention studies. Early positron emission tomography work reported right lateral cerebellar activation during silent recall of consciously retrieved episodic memories^20^. Subsequent task-based fMRI studies showed posterior cerebellar engagement during episodic retrieval, together with hippocampal and prefrontal activation^21^. More recent individual-level mapping has identified closely juxtaposed cerebellar territories corresponding to cerebral networks, revealing selective responses of cerebellar default mode network A during episodic projection^22^. Causal evidence is also emerging. Theta-frequency transcranial magnetic stimulation of the right cerebellum can enhance episodic encoding and later retrieval^23^, and right cerebellar transcranial direct-current stimulation has been reported to improve episodic memory in older adults both immediately and at six-month follow-up^24^. Together, these suggest that the cerebellum is not merely co-activated during memory tasks, but contribute to memory-relevant computations.

Despite these advances, a mechanistic account of cerebellar contribution to episodic memory in older adults remains incomplete. First, most evidence comes from younger adults, leaving uncertain whether the same cerebellar organization supports memory in ageing. Second, older-adult studies have often used relatively small samples, limiting capacity to capture consistent patterns across the population. Third, many studies have relied on resting-state fMRI, which maps intrinsic organization but cannot capture demand-dependent configurations. Fourth, it remains unclear whether the cerebellum supports memory through a unitary domain-general process or through multiple specialized subregions, each interacting with distinct cortical networks^18,25^. Finally, it is unknown whether cerebellar-cerebral interactions are best characterized by sustained coupling or by dynamic coordination that reconfigures across memory encoding and retrieval. This issue is particularly important in ageing, where reduced network segregation, neural dedifferentiation and altered temporal precision are commonly observed^6,26,27^.

Here, we examined task-evoked cerebellar organization and cerebellar–cerebral interactions during episodic memory in a large cohort of cognitively normal older adults with follow-ups (Fig. 1a-b). We addressed three questions. First, which cerebellar regions are recruited during episodic memory processing in older adults, and can task-responsive cerebellar activity be resolved into functionally dissociable networks with distinct behavioural relevance? To answer this, we applied sparse dictionary learning to task-activated cerebellar voxels and related network activities to recognition accuracy and response time (Fig. 1b-e). Second, how do memory-relevant cerebellar networks interact with cortical networks during encoding and retrieval? We quantified both static functional connectivity and dynamic functional connectivity to distinguish sustained coupling from memory-phase-dependent reconfiguration (Fig. 1f-g). Third, which patterns of cerebellar–cerebral interaction are behaviourally meaningful at baseline, and does the longitudinal stability of these dynamics predict subsequent memory change? We used dynamic time warping to compare connectivity trajectories across performance groups and tested whether trajectory stability predicted memory change over approximately four years (Fig. 1h-i). By placing the cerebellum within distributed memory circuits, this study extends models of memory ageing beyond a predominantly cortico-hippocampal framework.

**Fig. 1.**
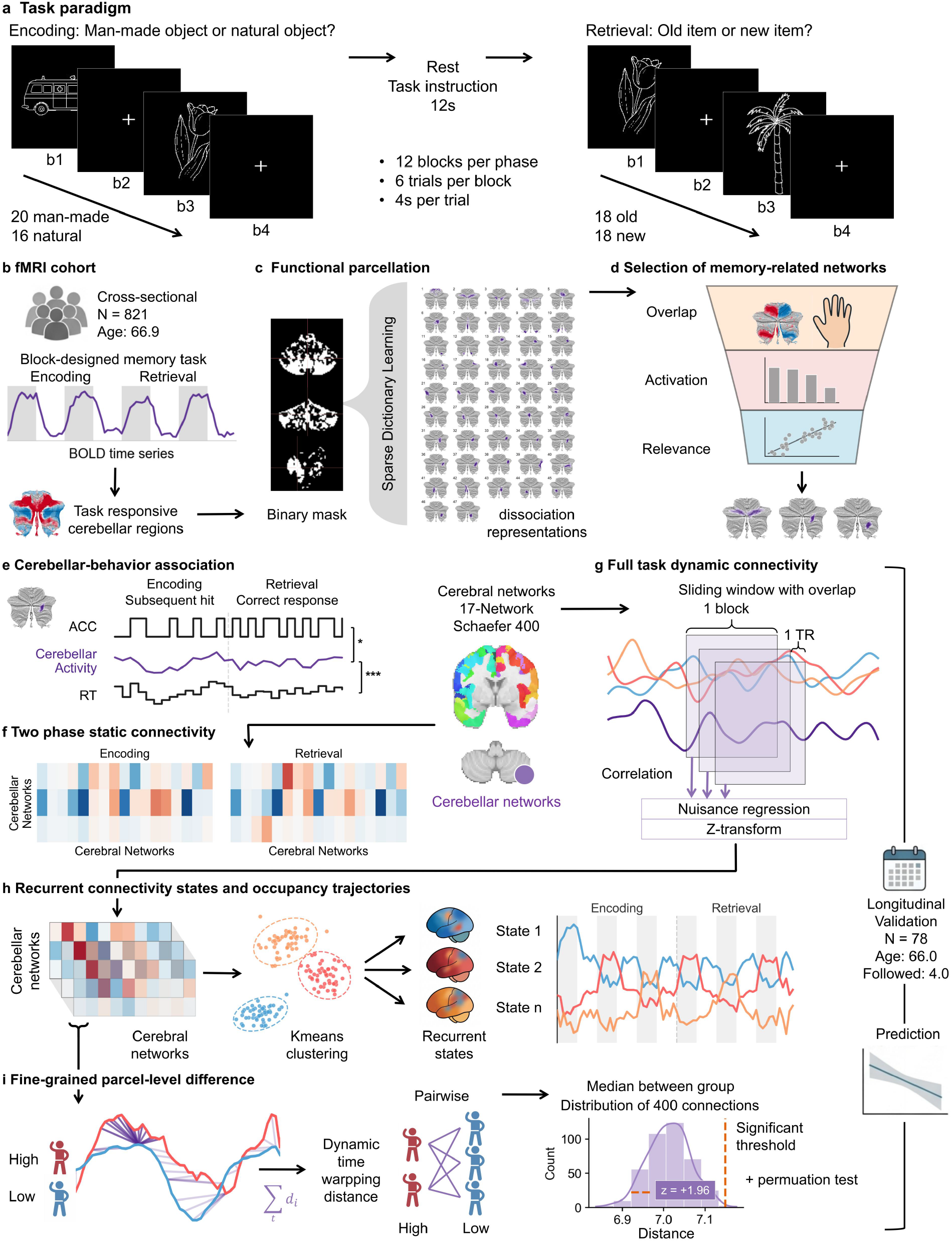
Research framework. **a**, Participants completed a block-designed episodic-memory fMRI task with separate encoding and retrieval phases. Each phase included 6 task blocks and 6 fixation blocks, with each block lasting 24 s. During encoding, participants viewed object images and judged whether each item was man-made or natural, a semantic task designed to promote incidental encoding. During retrieval, participants judged whether each item had appeared previously. Studied and novel items were presented in equal proportions, and the ratio of man-made to natural objects was matched to the encoding phase. **b**, The study included 821 cognitively normal older adults, 78 of whom had longitudinal follow-up data. Task contrasts were used to identify cerebellar regions engaged during memory processing. **c**, Task-responsive cerebellar regions were decomposed into dissociable functional networks using sparse dictionary learning. The cerebellar mask is displayed at x = 0, y = -60, and z = -30. **d**, Memory-relevant cerebellar networks were selected using a three-step procedure. Components were first excluded if more than 50% of their spatial extent overlapped with the left-versus-right motor-response contrast. Components showing significant activation in the motor-response contrast were then removed. Among the remaining non-motor components, networks significantly associated with task performance, including accuracy and response time, were retained and functionally characterized. **e**, For each memory-relevant cerebellar network, BOLD time series were extracted and related to behavioural time series during encoding and retrieval to characterize network-specific contributions to memory processing. **f**, Phase-specific static functional connectivity was computed between each memory-relevant cerebellar network and cortical parcels during encoding and retrieval. These analyses tested how cerebellar networks interacted with large-scale cortical systems and whether these interactions were associated with memory performance. Cortical regions were defined using the Schaefer 400-parcel atlas annotated by the Yeo 17-network solution. **g**, Dynamic functional connectivity was estimated during encoding and retrieval using sliding-window correlations. The window length matched one task block, and the step size was 1 TR. Raw connectivity time series were adjusted for covariates and z-standardized to obtain task-related dynamic connectivity trajectories. **h**, Recurrent connectivity states were identified from dynamic connectivity matrices using k-means clustering. This estimated recurrent connectivity patterns and their time-varying occupancy across the task. **i**, Fine-grained dynamic cerebellar–cerebral trajectories were analysed for each cerebellar network–cortical parcel pair. Dynamic time warping was used to quantify pairwise trajectory differences between participants from different performance groups. The median pairwise distance represented the between-group divergence for each connection. Significant trajectory differences were defined by two criteria: a distribution-based threshold exceeding 1.96 standard deviations and a permutation-test *P* value < 0.05. In the longitudinal subset, analyses in **e–i** were repeated to test how ageing altered these memory-related cerebellar mechanisms.

## Results

Participants completed a block-designed encoding-retrieval paradigm episodic memory task with separate encoding and retrieval phases (Fig. 1a). During encoding, they viewed line drawings of objects and made semantic category judgments to encourage semantic elaboration, indicating whether each item depicted a natural or man-made object. During retrieval, participants made old/new recognition judgments, discriminating previously studied items from novel foils.

### behavioural performance and longitudinal change

Participants showed high accuracy during encoding (0.94 ± 0.11), indicating adequate task engagement. Retrieval accuracy was 0.80 ± 0.12, with response time of 1,514 ± 341 *ms* Participants were more accurate when identifying new items than old, with new-item accuracy of 0.89 ± 0.14 and old-item accuracy of 0.70 ± 0.18. New-item judgments were also faster than old items, with response time of 1,467 ± 388 *ms* and 1,561 ± 357 *ms*, respectively. Signal detection analysis indicated moderate discrimination sensitivity, with *d*′ = 1.85 ± 0.69, and a positive response criterion of 0.45 ± 0.36 (Fig. 2a, Extended Data Table 1).

**Fig. 2.**
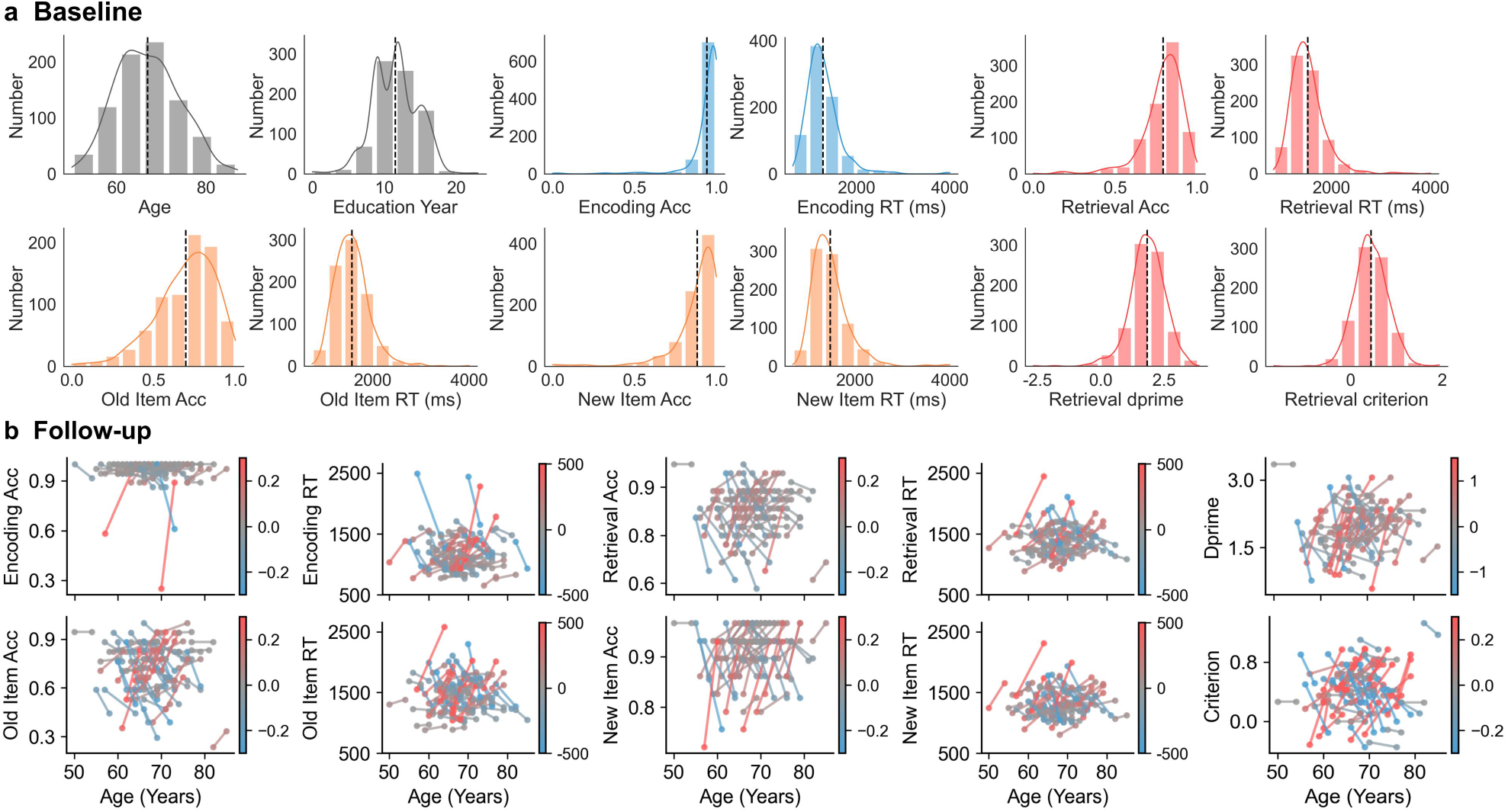
Task performance. **a**, Baseline demographic characteristics and task-performance distributions. Histograms show the distributions of age, sex, encoding and retrieval accuracy (Acc), response time (RT), old- and new-item accuracy and response time, retrieval discrimination sensitivity (dprime) and response criterion. The x axis indicates the value of each variable, and the y axis indicates the number of participants. **b**, Longitudinal changes in task performance across follow-up. The x axis indicates participant age, and the y axis indicates the value of each behavioural measure. Each line connects baseline and follow-up measurements for one participant. Red lines indicate an increase from baseline to follow-up, and blue lines indicate a decrease.

We stratified participants into three groups based on quartiles of retrieval accuracy. This stratification yielded 299, 271 and 251 participants in the high-, moderate- and low-performance groups, respectively (Extended Data Table 1). The three groups showed a graded profile across all measurements, supporting retrieval accuracy as a valid index to differentiate memory performance.

In the longitudinal subset, baseline task performance was comparable to the baseline. Encoding and retrieval accuracy was 0.96 ± 0.10 and 0.82 ± 0.08 at baseline. At follow-up, group-level performance remained stable, encoding and retrieval accuracy was 0.97 ± 0.05 and 0.83 ± 0.08 (Supplementary Table 1). Despite this group-level stability, individual changes were heterogeneous. Retrieval accuracy changed from −0.22 to 0.19, and retrieval response time changed from −609 to 950 *ms*. Item-specific accuracy changes ranged from −0.29 to 0.30 for old items and from −0.26 to 0.31 for new items (Fig. 2b). We further divided the longitudinal subset into memory maintainers (N = 48) and decliners (N = 30) according to the direction of change in retrieval accuracy (Supplementary Table 2-3).

### Episodic memory engaged widespread cerebellar regions in older adults

Voxel-wise task-activation analyses showed that both encoding and retrieval elicited widespread responses in bilateral posterior cerebellar lobules V–X with *P* <0.05 after false discovery rate (FDR) correction within the whole-brain mask (Fig. 3a). Positive task-related activation was observed in lobules V, VI, superior Crus I, VIIb, VIIIa, X. Negative activation was observed in inferior Crus I, Crus II and IX. Activation patterns for old and new item broadly resembled the retrieval map. The old-versus-new contrast showed effects in bilateral lobule V, superior VI, VI/Crus I boundary, clusters within Crus I, Crus II, VIIIa, and VIIIb. Motor-related activation was concentrated in lobules I–V, superior VI, VIII.

**Fig. 3.**
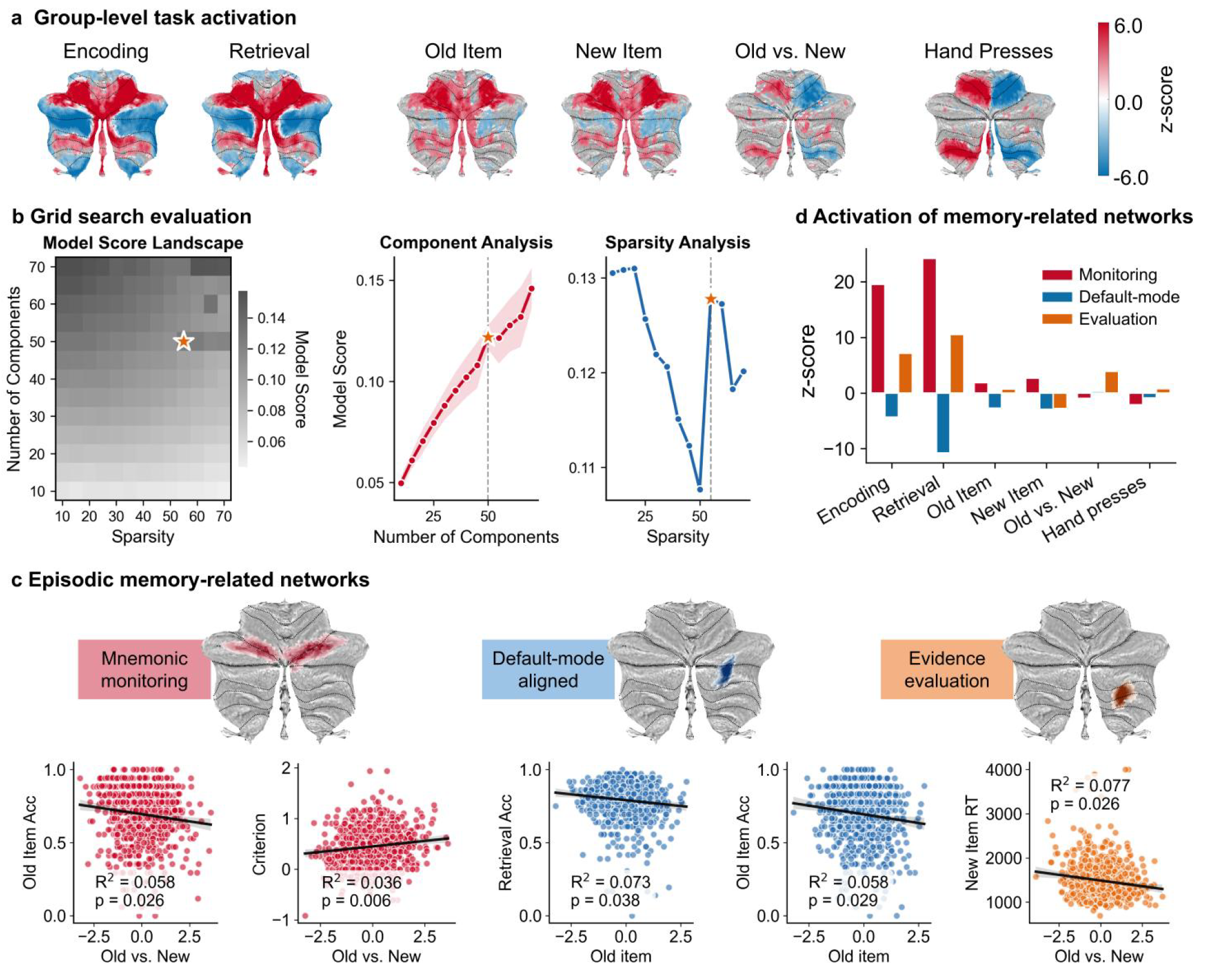
Task-responsive cerebellar networks and their associations with task performance. **a**, Group-level cerebellar activation across six task conditions. Red indicate positive task-evoked responses, and blue indicate negative responses. **b**, Data-driven decomposition of task-responsive cerebellar regions using sparse dictionary learning. The heat map summarizes model performance across the parameter space, with sparsity on the x axis, number of components on the y axis and colour denoting model score. Higher scores indicate better reconstruction. The line plots show model scores as a function of each parameter. In the component-number analysis, lines indicate scores averaged across sparsity values and shading denotes the standard deviation. The optimal solution comprised 50 components with a sparsity parameter of 55. **c**, Spatial maps and functional characterization of the three cerebellar networks significantly associated with task performance. In each scatter plot, the x axis shows the standardized activation estimate (z-scored) for the indicated task contrast, and the y axis shows the corresponding behavioural measure. Reported *P* values are Bonferroni-adjusted. Acc, accuracy; RT, response time.

No cerebellar voxels showed significant associations with task performance after FDR correction. This suggested the cerebellum may contribute through larger-scale functional networks rather than through isolated local responses, and motivated a network-level decomposition of task-responsive cerebellar activity.

### Data-driven decomposition identified three memory-relevant cerebellar networks

We applied sparse dictionary learning to task-responsive cerebellar voxels to identify functional networks without imposing anatomical boundaries. The optimal solution included a component number of 50 and a sparsity parameter of 55 according to elbow criterion of variance explained (Fig. 3b). The final solution retained 47 cerebellar networks after exclusion of brainstem components. These networks showed diverse bilateral, unilateral and vermal topographies, with over a half located in posterior regions (Extended Data Fig. 1).

To select memory-related networks, we excluded components with substantial motor overlap, and with significant hand response evoked activation, then related these non-motor networks’ activations to task performance. Five networks showed > 50% overlap with hand-presses activated region (Supplementary Table 4). Network-level general linear models (GLMs) showed other 7 networks were significantly activated by hand presses (Supplementary Fig. 1). Among the remaining non-motor components, 3 networks showed significant behavioural associations after Bonferroni correction, controlled for age, sex and years of education (Fig. 3c-d).

The first network, centred bilaterally at the lobule VI/Crus I boundary, was task-positive during both encoding and retrieval. Its old-versus-new activation was associated with old-item accuracy (*R*^*2*^ = 0.058, *β* = −0.020, *P* = 0.026) and response criterion (*R*^*2*^ = 0.036, *β* = 0.044, *P* = 0.006). We refer to this network as the mnemonic-monitoring network. The second network, localized to right Crus I/II, showed task-negative responses during both encoding and retrieval. Its old-item activation was associated with overall retrieval accuracy (*R*^*2*^ = 0.073, *β* = −0.016, *P* = 0.038) and old-item accuracy (*R*^*2*^ = 0.058, *β* = −0.024, *P* = 0.029). We refer to this network as the default-mode-aligned network. The third network, localized to right lobule VIIb/VIIIa, was task-positive during encoding and retrieval. This activation was associated with response time of new-item (*R*^*2*^ = 0.077, *β* = −43.125, *P* = 0.026). We refer to this network as the evidence-evaluation network.

All three networks showed stronger modulation during retrieval than during encoding, indicating preferential engagement during memory search and recognition decisions (Fig. 3d). Their spatial profiles were reproducible across model settings (Supplementary Fig. 2-4).

### Trial-level modulation linked cerebellar activity to response demand

To characterize how the three networks’ activities differed across performance groups, we examined their preprocessed, z-normalized BOLD time series and overlaid them onto trial-wise behavioural trajectories. We used haemodynamic response function (HRF)-informed modelling to test whether these activities varied with trial-level response accuracy and speed.

The mnemonic-monitoring network (Fig. 4a) showed the largest-amplitude, task-locked fluctuations among the three networks, with strong similarity across the high-, moderate-, and low-performance groups. Between-group correlations of the network time series were uniformly high during encoding (high versus moderate: *r* = 0.849, *P* < 0.001; high versus low: *r* = 0.846, *P* < 0.001; moderate versus low: *r* = 0.830, *P* < 0.001) and became even stronger during retrieval (high versus moderate: *r* = 0.882, *P* < 0.001; high versus low: *r* = 0.898, *P* < 0.001; moderate versus low: *r* = 0.937, *P* < 0.001). It showed positive response-time modulation during encoding across three groups (high: *β* = 0.035, *P* < 0.001; moderate: *β* = 0.025, *P* < 0.001; low: *β* = 0.038, *P* < 0.001), and during retrieval only in high performers (*β* = 0.025, *P* < 0.001).

**Fig. 4.**
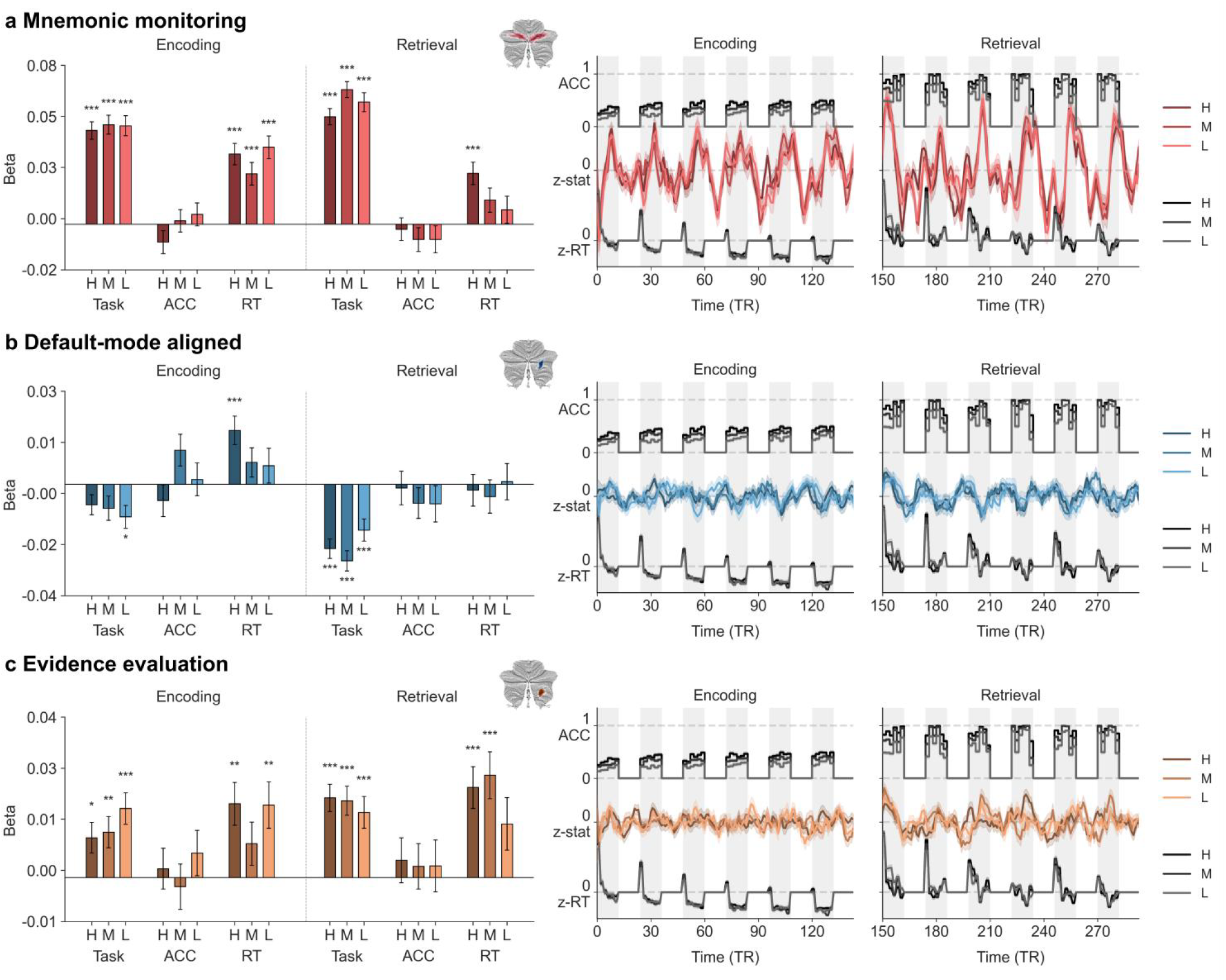
Temporal brain–behaviour coupling of memory-relevant cerebellar networks. Trial-level associations between cerebellar network activity and behavioural responses were estimated using HRF-informed models. Participants were stratified into high- (H), moderate- (M) and low-performance (L) groups. Model regressors included task events (task), trial-level accuracy (ACC) and response time (RT). During encoding, ACC denotes subsequent hit status; because encoded items appeared randomly during retrieval, group-level mean values range from 0 to 0.5. During retrieval, ACC denotes whether the current trial was answered correctly, with group-level mean values ranging from 0 to 1. For each cerebellar network, the left bar plots show beta estimates from the HRF-informed model and their statistical significance. Significance levels are indicated as ^***^*P* < 0.001, \*\**P* < 0.01 and ^*^*P* < 0.05. The right panels show group-averaged cerebellar activity time series and behavioural response time series. For visualization, brain activity time series were smoothed with a 6-TR window; shading denotes the standard deviation. Colour intensity indicates memory-performance group, with darker, intermediate and lighter colors representing high-, moderate- and low-performance groups, respectively.

The default-mode-aligned network (Fig. 4b) showed modest inter-group similarity during encoding (high versus moderate: *r* = 0.291, *P* < 0.001; high versus low: *r* = 0.167, *P* < 0.001; moderate versus low: *r* = 0.457, *P* < 0.001). During retrieval, however, the temporal profiles became more aligned across groups (high versus moderate: *P* = 0.522, *P* < 0.001; high versus low: *r* = 0.549, *P* < 0.001; moderate versus low: *r* = 0.522, *P* < 0.001). It showed sustained negative retrieval-phase-related modulation, but response-time modulation was largely absent except during encoding in high performers (*β* = 0.017, *P* < 0.001).

The evidence-evaluation network (Fig. 4c) showed relatively low-amplitude oscillations. During encoding, inter-group similarity ranged from modest to moderate (high versus moderate: *r* = 0.156, *P* = 0.061; high versus low: *r* = 0.350, *P* < 0.001; moderate versus low: *r* = 0.576, *P* < 0.001), whereas during retrieval similarity increased (high versus moderate: *r* = 0.393, *P* < 0.001; high versus low: *r* = 0.516, *P* < 0.001; moderate versus low: *r* = 0.615, *P* < 0.001). It showed the most consistent response-time effects across encoding (high: *β* = 0.017, *P* = 0.002; moderate: *β* = 0.008, *P* = 0.206; low: *β* = 0.017, *P* = 0.005) and retrieval (high: *β* = 0.021, *P* < 0.001; moderate: *β* = 0.024, *P* < 0.001; low: *β* = 0.012, *P* = 0.08).

These indicated that temporal cerebellar activities were closely related to processing demand and decision efficiency. Across memory-performance groups, task-evoked activities were most similar in the mnemonic-monitoring network, followed by the default-mode-aligned network, and then the evidence-evaluation network, with greater between-group similarity during retrieval than during encoding for all of the three.

### Static connectivity revealed stable cerebellar–cerebral coupling

Having identified the three memory-related cerebellar networks and characterized their contributions, we next examined their phase-specific static functional connectivities with the cerebral cortex. For each network, connectivity was estimated across all time points within encoding and retrieval phases to characterize sustained coupling patterns. Each network showed a distinct but highly consistent connectivity profile across two phases (Fig. 5a).

**Fig. 5.**
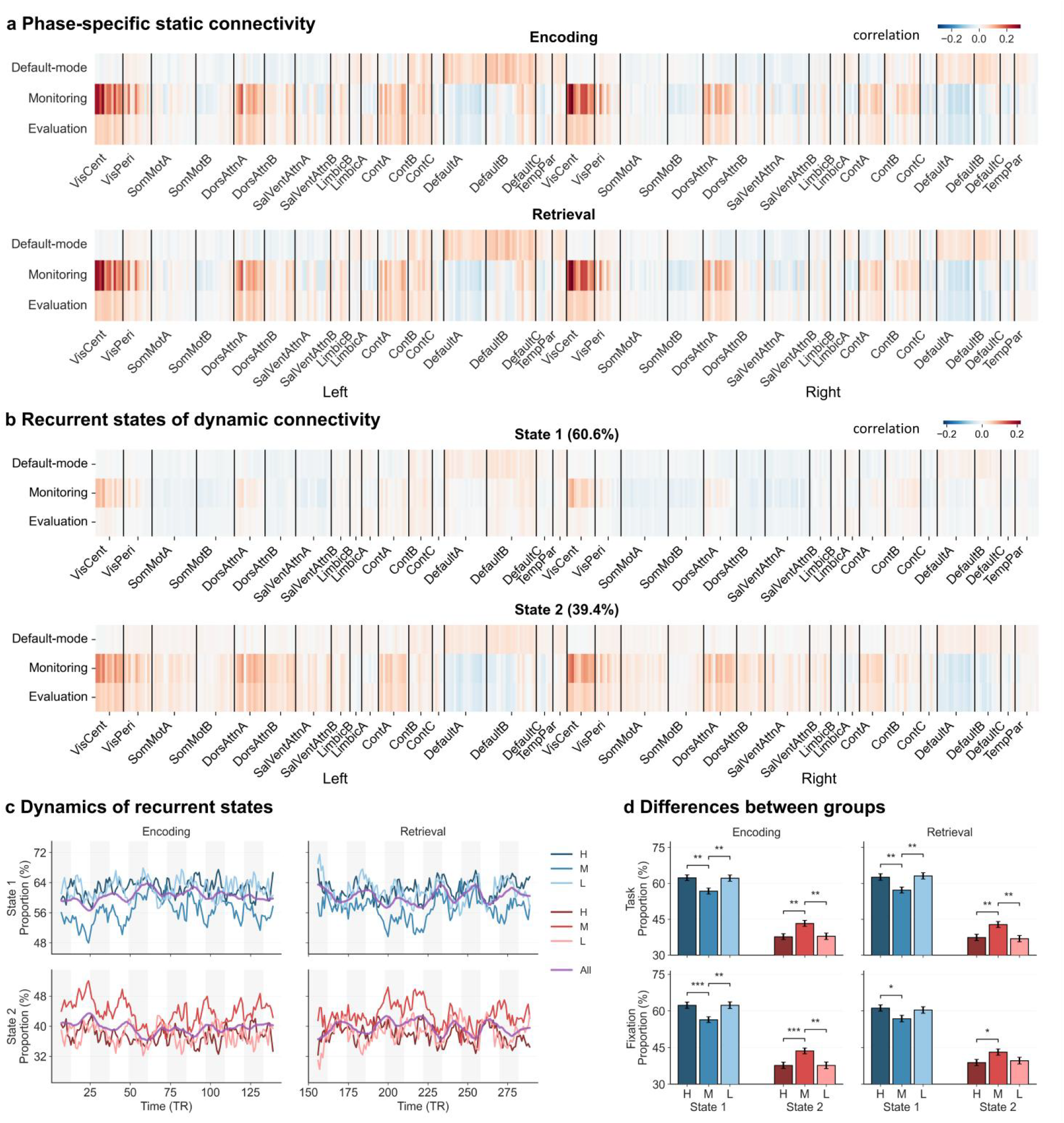
Cerebellar–cerebral functional connectivity profiles during the task. **a**, Phase-specific static functional connectivity between the three memory-relevant cerebellar networks and cortical parcels during encoding (top) and retrieval (bottom). The x axis shows 400 cortical parcels grouped by the Yeo 17-network solution and separated by hemisphere. The y axis shows the three cerebellar networks. **b**, Recurrent cerebellar–cerebral connectivity states derived from dynamic functional connectivity time series. State 1 (top), the dominant task-related state, was characterized by widespread weak negative connectivity. State 2 (bottom), the less frequent state, showed a connectivity pattern more similar to the phase-averaged static functional connectivity profiles. **c**, Time-varying occupancy of the two recurrent states across the task. State 1 (top, blue) increased during task blocks, whereas State 2 (bottom, red) showed a complementary pattern. Each time point represents the centre of the sliding window used to estimate dynamic functional connectivity, with the value calculated from data spanning half a block before and half a block after that point. Colour intensity indicates memory-performance group, with darker, intermediate and lighter colors representing high-, moderate- and low-performance groups, respectively. Purple lines show the group-level average across all participants. **d**, Mean occupancy of State 1 and State 2 during task blocks (top) and fixation blocks (bottom) in the high-, moderate- and low-performance groups. Significance indicates between-group differences in state occupancy. Significance levels are indicated as \*\*\**P* < 0.001, \*\**P* < 0.01 and \**P* < 0.05.

The default-mode-aligned network is mainly positively connected with the cerebral default mode networks. The strongest coupling was with default-B, followed by default-A, default-C, and temporoparietal networks, concentrated in prefrontal and temporal parcels. Negative connectivities were most evident with dorsal-attention-A, and salience/ventral-attention networks. The mnemonic-monitoring network showed strongest positive connectivity with central and peripheral visual network during both encoding and retrieval, concentrated in bilateral extrastriate visual parcels. It also showed positive coupling with dorsal-attention-A, control-A, contol B networks. Negative coupling was observed with default-A and limbic-A networks. The evidence-evaluation network was also positively connected with visual, dorsal-attention-A, and control networks, with the strongest parcels in bilateral extrastriate, superior parietal and intraparietal/frontoparietal regions, and negatively connected with default-A network. It showed extra positive coupling with limbic-A networks.

These static connectivity profiles were largely preserved across encoding and retrieval, but did not differ significantly across memory-performance groups after Bonferroni correction.

### Recurrent dynamic states of dynamic connectivity captured task demands

To capture the temporal evolution of task-related connectivity, we calculated sliding-window cerebellar–cerebral dynamic functional connectivity, and used K-means clustering to identify recurrent connectivity states in the task. A two-state solution provided the best fit, yielding the highest silhouette coefficient among solutions with k = 2–5 (Supplementary Fig. 5). State 1 reflected a relatively weak configuration, with limited or negative coupling to sensorimotor, dorsal-attention-B and salience-attention-A networks. State 2 reflected stronger and more segregated configuration, similar to the static connectiity (Fig. 5b).

State expression fluctuated across the task (Fig. 5c). State 1 was predominant throughout the task, comprising approximately 48–67%. Its occupancy increased during task blocks and attenuated during fixation, with the reciprocal pattern for State 2. Similar task-locked fluctuations were observed during retrieval. This indicated that episodic memory processing demand shifted cerebellar–cerebral dynamics toward State 1, with reversion toward State 2 during fixation. Performance groups differed in state occupancy during both encoding and retrieval, for both task and fixation blocks (Fig. 5d). High and low performers spent more time in State 1 than moderate performers, particularly in fixation blocks during encoding.

### Parcel-level cerebellar–cerebral dynamic trajectories differentiated memory-performance groups

We next examined whether the divergence of state occupancy was expressed in spatially fine-grained parcel-level connections. For each cerebellar network – cerebral parcel pair, we used dynamic time warping distance (*d*) to quantify between-group difference in dynamic connectivity trajectories. We found that performance-related trajectory differences were more pronounced during retrieval than encoding and were distributed across cortical systems in a network-specific manner.

During encoding (Fig. 6a), only the default-mode-aligned network showed significant difference. The comparison between the high- and moderate-performing groups revealed a significant difference on the left ventral prefrontal default-B network (*d* = 7.764, *Z* = 2.919, *P* = 0.047). The moderate-versus-low differences involved the left superior parietal dorsal-attention-A network (*d* = 7.647, *Z* = 2.018, *P* = 0.010), and the right inferior parietal default-C network (*d* = 7.644, *Z* = 1.965, *P* = 0.028). The default-mode-aligned network was also involved during retrieval (Fig. 6b), with significant differences observed in the left ventral prefrontal default-B network, and the right somatomotor-A network (*d* = 7.051, *Z* = 2.056, *P* = 0.006) between high and moderate performers (*d* = 7.062, *Z* = 2.262, *P* = 0.037). The moderate-versus-low differences involved the left temporo-occipital dorsal-attention-A network (*d* = 7.177, *Z* = 2.233, *P* = 0.027).

**Fig. 6.**
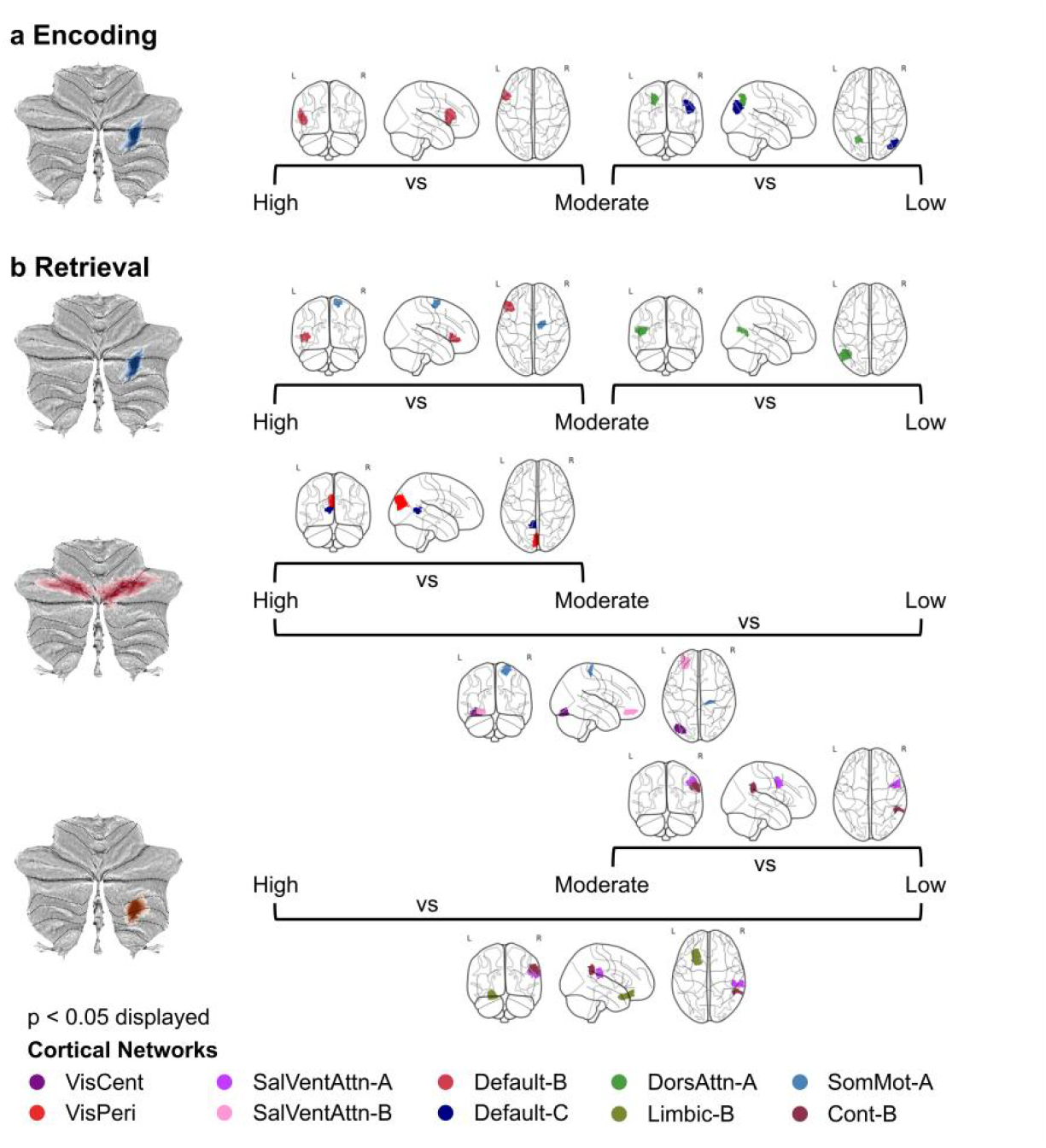
Between-group differences in dynamic cerebellar–cerebral connectivity trajectories. Significant cerebellar–cerebral network pairs are shown for task phases in which dynamic functional connectivity trajectories differed between performance groups. The cerebellar network is shown on the left, and the cortical parcel showing a significant between-group difference in pairwise dynamic connectivity is shown on the right. Significant effects met both criteria: a standardized distributional difference > 1.96 and a permutation-test *P* < 0.05 based on 5,000 permutations. **a**, During encoding, significant trajectory differences were observed only for the default-mode-aligned network. **b**, During retrieval, all three cerebellar networks showed significant trajectory differences with specific cortical parcels. VisCent, central visual network; VisPeri, peripheral visual network; SalVentAttn, salience/ventral-attention network; Default, default-mode network; DorsAttn, dorsal-attention network; Limbic, limbic network; SomMot, somatomotor network; Cont, control network.

The mnemonic-monitoring network differentiated performance groups mainly during retrieval. In the high-versus-low comparison, significant trajectory differences were observed for the left insular salience/ventral-attention-B network (*d* = 7.094, *Z* = 2.054, *P* = 0.004), the right somatomotor-A network (*d* = 7.095, *Z* = 2.066, *P* = 0.017), and the left extrastriate peripheral visual network (*d* = 7.091, *Z* = 1.997, *P* = 0.031). The high-versus-moderate differences involved the left superior extrastriate central visual network (*d* = 7.096, *Z* = 2.764, *P* = 0.025), and the left retrosplenial default-C network (*d* = 7.055, *Z* = 2.014, *P* < 0.001).

Differences were also prominent for the evidence-evaluation network during retrieval. The high-versus-low differences involved the right temporal control-B network (*d* = 7.108, *Z* = 2.236, *P* = 0.010), the right frontal-eye-field dorsal-attention-A network (*d* = 7.099, *Z* = 2.068, *P* = 0.042), and the left orbitofrontal limbic-B network (*d* = 7.095, *Z* = 2.011, *P* = 0.011). In the moderate-versus-low comparison, differences involved the right parietal opercular salience/ventral-attention-A network (*d* = 7.181, *Z* = 2.664, *P* = 0.024), and the right temporal control network (*d* = 7.166, *Z* = 2.385, *P* = 0.027).

Thus, differences in episodic memory performance were not primarily reflected on static functional connectivity. Instead, they were associated with the temporal dynamics and alignment of specific cerebellar–cerebral interactions.

### In longitudinal subset, cerebellar activations were preserved, but activity-performance associations were divergent

In longitudinal subest, we investigated whether changes in memory performance were associated with altered activation of the three memory-relevant cerebellar networks, and whether these alterations distinguished maintainers from decliners. We found that group-level activation at follow-up broadly recapitulated the baseline patterns. Despite the overall stability, among maintainers, old-versus-new modulation increased significantly in the default-mode-aligned network but decreased in the evidence-evaluation network. Among decliners, old-versus-new modulation increased differently in the mnemonic-monitoring network (Fig. 7a).

**Fig. 7.**
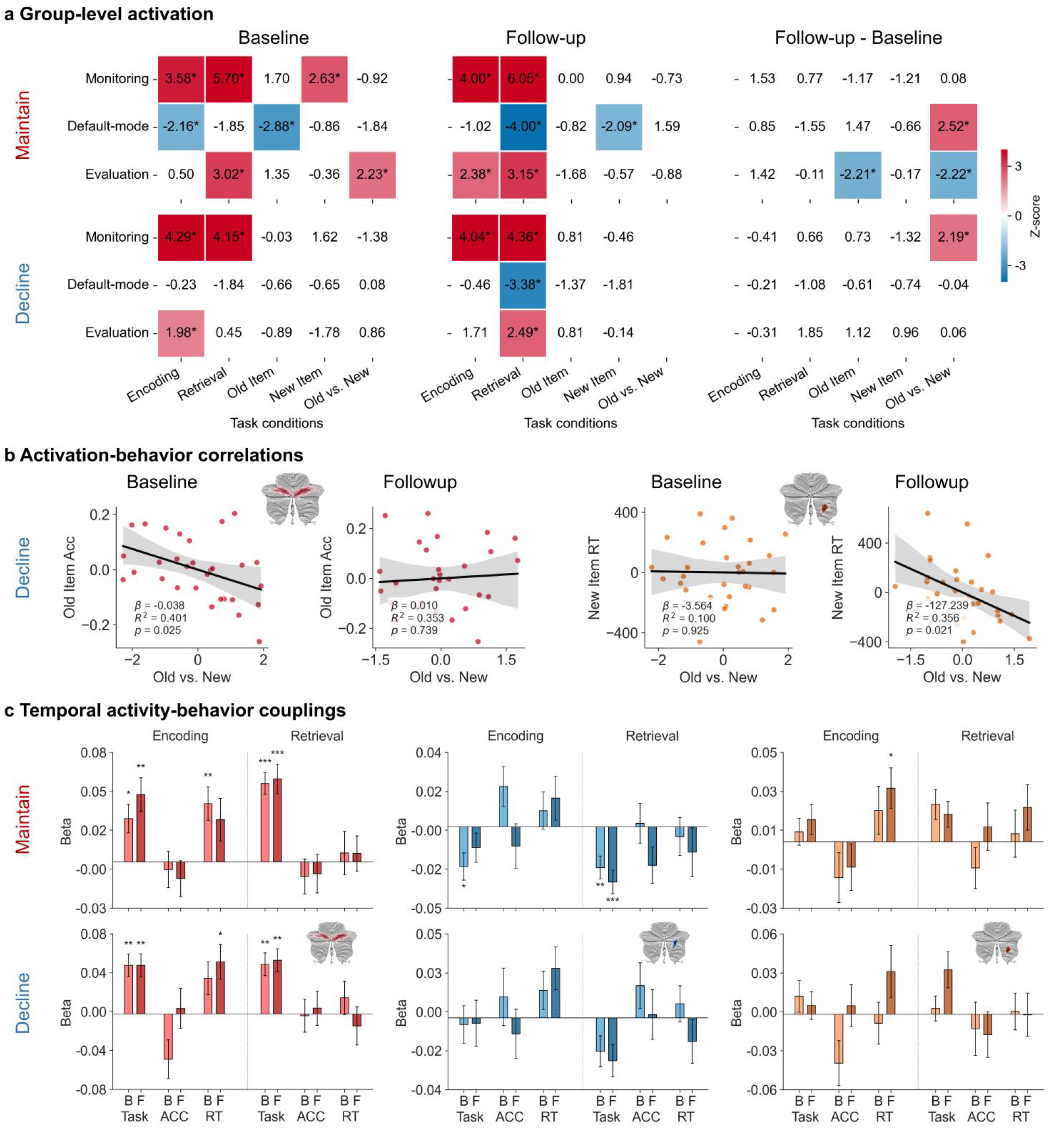
Longitudinal cerebellar activity in memory maintainers and decliners. Participants with longitudinal follow-up were classified as memory maintainers or decliners to examine neural mechanisms associated with memory resilience and decline. **a**, Group-level activation profiles of the three memory-relevant cerebellar networks in maintainers (top) and decliners (bottom). Activation is shown at baseline (left), follow-up (middle) and for the follow-up-minus-baseline contrast (right). In each heat map, the x axis indicates task contrast and the y axis indicates cerebellar network. Warm colors indicate positive activation, and cool colors indicate negative activation. ^*^*P* < 0.05 after FDR correction. **b**, Changes in activation–behaviour associations in decliners. For the mnemonic-monitoring network (left two scatter plots), significant negative associations observed at baseline were attenuated at follow-up. For the evidence-evaluation network (right two scatter plots), significant negative associations emerged at follow-up. The x axis shows the standardized activation estimate (z score) of the corresponding cerebellar network under the indicated task condition, and the y axis shows task performance, including accuracy (Acc) or response time (RT). **c**, Trial-level associations between cerebellar network activity and task events or behavioural responses in maintainers (top) and decliners (bottom). Results are shown for the mnemonic-monitoring network (left), default-mode-aligned network (middle) and evidence-evaluation network (right). Associations were estimated using HRF-informed models with regressors for task events, trial-level accuracy (ACC) and response time (RT). During encoding, ACC denotes subsequent hit status. During retrieval, ACC denotes whether the current trial was answered correctly. Significance levels are indicated as ^***^*P* < 0.001, ^**^*P* < 0.01 and ^*^*P* < 0.05.

Changes in activation–performance associations were observed only among decliners. At baseline, stronger old-versus-new activation in the mnemonic-monitoring network was associated with lower accuracy for old items, but this association was absent at follow-up. Instead, stronger old-versus-new activation in the evidence-evaluation network at follow-up was associated with faster responses to new items (Fig. 7b).

In trial-level models, among maintainers, the modulation of encoding activity by response time weakened in the mnemonic-monitoring network but emerged in the evidence-evaluation network at follow-up. Task-negative retrieval activity in the default-mode-aligned network remained stable. Among decliners, the association between mnemonic-monitoring activity and encoding response time strengthened, whereas the default-mode-aligned network showed little relationship with task structure at either time point (Fig. 7c).

### Static connectivity remained stable, whereas recurrent state occupancy changed

Longitudinal analyses of static connectivity revealed no significant changes in either group. Projecting baseline and follow-up dynamic connectivity data onto the two previously identified recurrent states showed that their overall balance and task-evoked fluctuations were preserved over time. However, state occupancy changed among decliners: during encoding blocks, State 1 occupancy increased, whereas State 2 occupancy decreased (Fig. 8a, left). Analyses of state dynamics during encoding and retrieval showed stable temporal patterns in maintainers but marked changes in decliners, particularly during retrieval (Fig. 8a, right). Thus, longitudinal memory decline was associated with altered temporal deployment of pre-existing cerebellar– cerebral connectivity states.

**Fig. 8.**
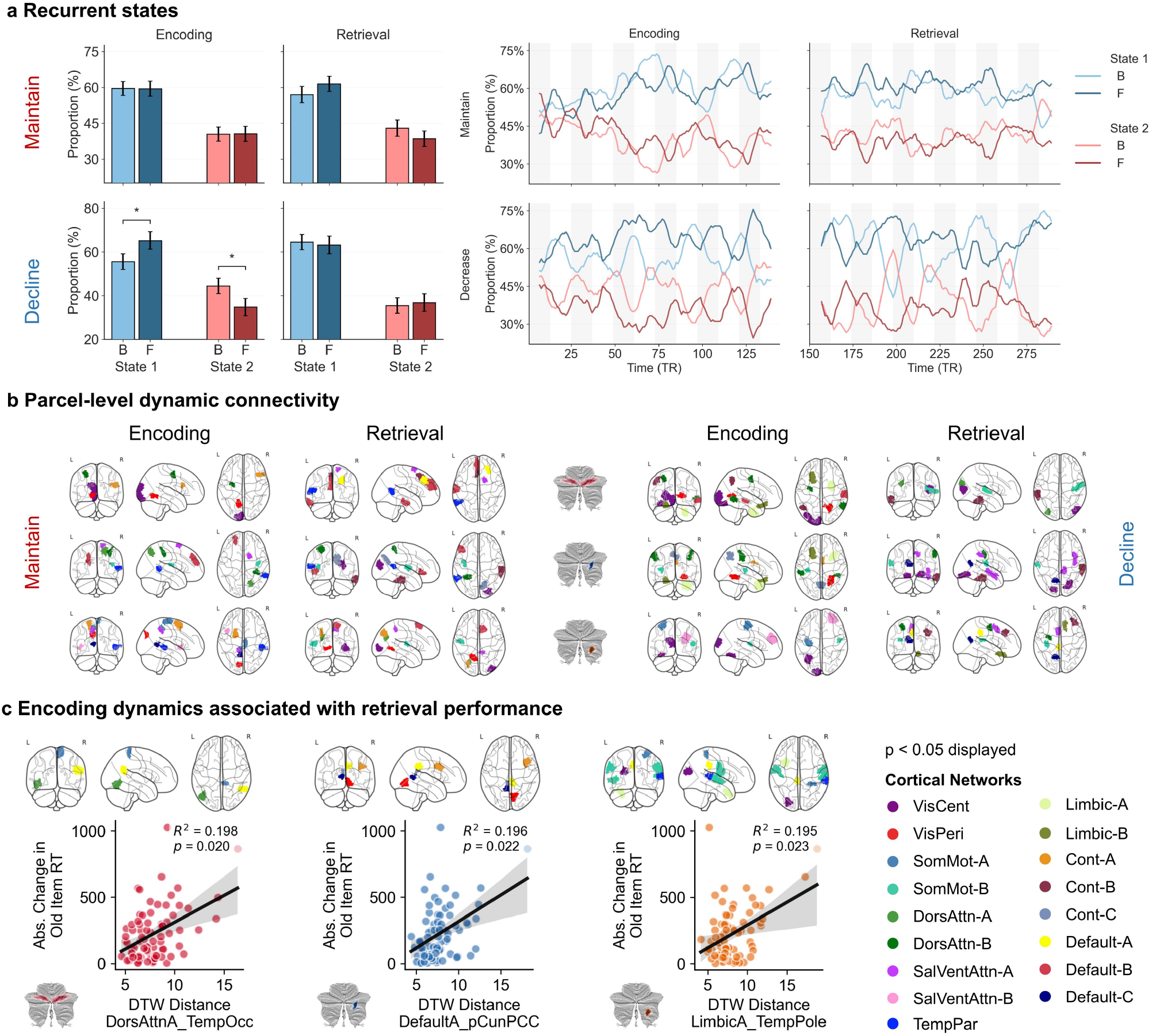
Longitudinal changes in cerebellar-cerebral functional connectivity and their association with memory performance change. **a**, Longitudinal changes in recurrent dynamic-state occupancy. The left bar plots show the mean occupancy of State 1 (blue) and State 2 (red) during task blocks in memory maintainers (top) and decliners (bottom). Significance levels are indicated as *\*P* < 0.05. The right panels show the time-varying occupancy of each state across the task. Light colors indicate baseline and dark colors indicate follow-up. Decliners showed increased State 1 occupancy during encoding. **b**, Longitudinal differences in dynamic cerebellar–cerebral connectivity trajectories. Significant cerebellar–cerebral network pairs are shown for task phases in which dynamic functional connectivity trajectories differed between baseline and follow-up. Results are shown separately for maintainers (left) and decliners (right). Each row shows one cerebellar network and the cortical parcels with which it exhibited significant longitudinal trajectory differences. Trajectory differences involving the temporoparietal network were observed only in maintainers, whereas differences involving the limbic network were observed only in decliners. **c**, Associations between longitudinal instability of encoding-phase dynamic connectivity and changes in retrieval response time. Dynamic time warping (DTW) distance between baseline and follow-up connectivity trajectories significantly predicted longitudinal changes in task response time, especially for old-item response time. The left, middle and right panels show predictive cerebellar-cerebral connections for the mnemonic-monitoring, default-mode-aligned and evidence-evaluation networks, respectively. The number of predictive connections increased from the mnemonic-monitoring network to the default-mode-aligned network and was greatest for the evidence-evaluation network. For each cerebellar network, one representative scatter plot shows the association between DTW distance for a specific cerebellar network–cortical parcel pair and the absolute longitudinal change in old-item response time. The x axis indicates DTW distance, and the y axis indicates the absolute change in old-item RT. VisCent, central visual network; VisPeri, peripheral visual network; SomMot, somatomotor network; DorsAttn, dorsal-attention network; SalVentAttn, salience/ventral-attention network; TempPar, temporoparietal network; Limbic, limbic network; Cont, control network; Default, default-mode network. Suffixes A, B and C indicate corresponding subnetworks.

### Parcel-level dynamics differentiated memory ageing and predicted retrieval efficiency

We next examined whether longitudinal instability in fine-grained cerebellar– cerebral coupling was associated with memory change. For each cerebellar network– cortical parcel pair, we calculated the dynamic time-warping distance between baseline and follow-up trajectories, larger distances indicated greater instability. Longitudinal changes were group-specific: altered coupling with the temporoparietal network in maintainers, with the limbic-A and limbic-B networks in decliners (Fig. 8b). During encoding, changes in maintainers involved temporoparietal network and each of the evaluation and default-mode networks. In decliners, altered coupling involved the monitoring and default-mode networks in relation to the limbic-A and B networks. During retrieval, maintainers showed altered coupling between the monitoring network and temporoparietal network, whereas decliners showed altered coupling between the evaluation network and limbic-B network.

Finally, we tested whether trajectory instability predicted continuous changes of memory performance. After Bonferroni correction, greater instability during encoding robustly predicted longitudinal retrieval slowing, particularly for old-items (Fig. 8c). For the mnemonic-monitoring network, greater encoding-phase instability with the left temporo-occipital dorsal-attention network (*R*^*2*^ = 0.198, *P* = 0.020) and the right somatomotor-A network (*R*^*2*^ = 0.179, *P* = 0.049) predicted old-item slowing. Instability with the right inferior parietal default-A network predicted overall retrieval slowing (*R*^*2*^ = 0.194, *P* = 0.038). For the default-mode-aligned network, greater instability with the right posterior cingulate/precuneus default-A network (*R*^*2*^ = 0.196, *P* = 0.022), right lateral prefrontal control-A network (*R*^*2*^ = 0.181, *P* = 0.045), and left retrosplenial default-C network (*R*^*2*^ = 0.180, *P* = 0.048) predicted old-item slowing.

Instability with the right peripheral visual network (*R*^*2*^ = 0.201, *P* = 0.026) and left posterior cingulate control-C network (*R*^*2*^ = 0.191, *P* = 0.043) predicted overall retrieval slowing. The evidence-evaluation network showed the most extensive effects, predominantly through encoding-phase coupling with somatomotor, visual, limbic, default-mode, and temporoparietal networks. The strongest association with old-item response-time slowing was observed for the left auditory somatomotor-B network (*R*^*2*^ = 0.259, *P* < 0.001). Additional predictors of old-item slowing included the left temporal-pole limbic (*R*^*2*^ = 0.195, *P* = 0.023), the left extrastriate visual network (*R*^*2*^ = 0.188, *P* = 0.033), and the left posterior cingulate/precuneus default-A networks (*R*^*2*^ = 0.182, *P* = 0.043). Instability with the right temporoparietal networks (*R*^*2*^ = 0.192, *P* = 0.042) predicted overall retrieval slowing.

Together, these reveal a hierarchy of sensitivity in memory ageing: activation and static connectivity were largely preserved, whereas recurrent state occupancy captured performance change. Parcel-level dynamic connectivity further predicted retrieval slowing.

## Discussion

In a large cohort of older adults, we showed that the cerebellum contributed to episodic memory through specialized functional networks and their dynamic coordination with cerebral networks. Although task-evoked cerebellar responses were widespread, individual voxels showed little behavioural specificity. Data-driven decomposition instead revealed three non-motor networks with distinct response profiles, cortical affiliations and behavioural correlates: a bilateral lobule VI/Crus I mnemonic-monitoring network associated with old-item recognition accuracy and response criterion; a right Crus I/II default-mode-aligned network associated with recognition accuracy, particularly for old items; and a right lobule VIIb/VIIIa evidence-evaluation network associated with response time. Temporally resolved, parcel-level cerebellar–cerebral coordination explained individual differences in memory performance and longitudinal changes in retrieval efficiency better than mean activation, phase-specific static connectivity or coarse recurrent states. These findings extend models of memory ageing beyond cortico-hippocampal circuitry by placing posterior cerebellar networks within the distributed systems that support encoding, recognition and memory-guided decisions.

The cerebellum should therefore be viewed not as a unitary auxiliary structure recruited during memory processing, but as a collection of specialized networks that support distinct components of episodic memory. The mnemonic-monitoring network was coupled with visual, attention, and control networks, aligning it with visual– frontoparietal systems preferentially engaged by novel or externally driven information^28^. During retrieval, differences in its dynamic connectivity across performance groups were most pronounced in the central visual, default-mode and salience networks. This profile accords with theories proposing that the cerebellum supports temporal prediction, error monitoring and adaptive calibration of cortical processing^17,18^. The default-mode-aligned network was coupled with medial prefrontal, temporal and posterior default-mode regions. This connectivity links it to systems that preferentially process familiar or previously encountered information^28^ and support internally oriented cognition, autobiographical construction and memory-guided simulation^6,7^. The evidence-evaluation network was coupled with visual, attention, control, and limbic networks. This broad connectivity reflects the multiple demands of recognition decisions, including sensory analysis, attentional allocation, comparison with stored representations and response selection^29^. Comparisons with published cerebellar atlases further supported these functional profiles. The mnemonic-monitoring, default-mode-aligned and evidence-evaluation networks overlapped most strongly with visual–attentional, default-mode and control– attentional territories, respectively (Supplementary Fig. 6). These profiles accord with contemporary accounts of cerebellar topography, which identify multiple cognitive territories within Crus I/II and adjacent posterior lobules^12,13,25,30^. These interpretations of the three specialized networks accord with cerebellar theories emphasizing predictive timing, sequencing, and adaptive updating^16,17,31^ and extends them to episodic-memory in older adults.

A central implication is that cerebellar contributions to episodic memory operate at fine temporal and spatial scales. The three cerebellar networks coordinated with cortical systems according to task phase and memory performance, consistent with evidence for segregated cerebellar–cerebral loops involving prefrontal, parietal, and temporal cortices^9,10^. Episodic memory involves temporally ordered processes, including visual perception, sustained attention, semantic elaboration, memory formation, memory search, and evidence evaluation^32,33^. Activation and connectivity averaged across an entire task phase may obscure this time-sensitive coordination. In our study, static connectivity revealed a clear network architecture, but remained largely stable across encoding and retrieval and showed limited behavioural relevance. Recurrent dynamic states captured task context, with one increased its occupancy during task blocks, and the other during fixation. More importantly, fine-grained dynamic trajectories between distributed cerebellar networks and cortical parcels differed across performance groups. Thus, memory performance may depend not simply on whether cerebellar and cortical networks are connected, but on whether they interact in the appropriate regions at the appropriate times^34^. The longitudinal findings validate this interpretation and further reveal how specialized cerebellar networks change their activities during memory ageing. Mean cerebellar activation remained largely stable, suggesting they remain recruitable in cognitively normal older adults. However, instability of encoding-phase dynamic coupling predicted later retrieval slowing, especially for old-item recognition. Overall, individual and longitudinal differences in older adults reflect not only on averaged activation or connectivity strength but also the temporal alignment of specialized cerebellar networks with specific cortical processing streams.

The longitudinal findings on cerebellar networks are consistent with features of cognitive ageing: neural dedifferentiation, reduced network segregation, and impaired temporal precision^26,27^. First, the altered activation-behaviour associations suggest neural dedifferentiation: in decliners, mnemonic-monitoring activation was stronger, but it was no longer associated with old-item recognition as it had been at baseline, indicating broader but less functionally specific recruitment. Second, reduced network segregation was evident in decliners, whose longitudinal change was mainly expressed in State 1. This task-related recurrent state showed widespread weak negative coupling and lower modularity than the more segregated fixation-related configuration. In maintainers, by contrast, State 1 occupancy remained stable over time. Third, impaired temporal precision was evident in the dynamic occupancy of State 1: decliners showed much larger differences across the two follow-ups, whereas maintainers remained relatively stable, suggesting that memory support depends on recruiting the state at the right time.

Methodologically, these findings indicate that network-level and time-resolved analyses are well suited to the cerebellum’s distributed and dynamic functional organization. The absence of robust voxel-wise behavioural associations, together with the emergence of network-level effects, suggests that dimensionality-reduction approaches may capture cerebellar function more effectively than isolated voxel-wise analyses. This interpretation is consistent with previous work showing that cerebellar contributions to memory become more apparent at the network level^35^. A key strength of the present analysis is that the networks were derived from task-responsive cerebellar voxels without imposing predefined lobular or atlas-based boundaries. This approach is important because cerebellar functional fields are spatially compact, partially overlapping, and often poorly represented by coarse anatomical divisions^22,36,37^. The resulting components indicate that Crus I/II does not constitute a homogeneous cognitive region. Instead, adjacent posterior territories supported dissociable memory-related processes: the lobule VI–Crus I boundary was associated with visual–attentional monitoring, Crus I/II was embedded in default-mode circuitry, and the lobule VIIb/VIIIa network was coupled to attentional, control, and limbic systems. This functional fractionation resembles previous descriptions of hierarchical cognitive control in the cerebellum^38^, and extends them to episodic memory. It also provides a network-level explanation for why previous imaging and stimulation studies have implicated the cerebellum in both episodic encoding and retrieval, and why cerebellar effects vary across task types, memory phases, and behavioural measures.

Several limitations should be noted. First, the task involved only item recognition. Future studies should examine whether similar cerebellar dynamics support more naturalistic and ecologically valid forms of memory, including associative learning, spatial navigation, and autobiographical recollection. Second, the used fMRI data has limited temporal resolution and cannot fully capture the rapid dynamics of cerebellar activity or cerebellar–cerebral interactions. Combining ultrafast fMRI and other high-temporal-resolution methods could characterize these interactions more precisely. Third, dynamic connectivity and dynamic time warping quantify temporal coordination but do not establish the direction of information flow or causal influence. Future studies should combine directional connectivity models, perturbational methods, and cerebellar stimulation to test these causal mechanisms. Fourth, the longitudinal subset was relatively small. Replication in larger longitudinal cohorts and independent datasets is needed. Finally, we observed heterogeneity in cerebellar activity within the same performance. Precision mapping may help explain this heterogeneity and clarify how individual-specific cerebellar activities contribute to episodic memory^14,39^.

## Conclusion

This study identified the cerebellum as a functionally specialized and dynamically coordinated component of episodic memory in older adults. Task-responsive cerebellar regions were organized into specialized functional networks implicated in mnemonic-monitoring, default-mode-aligned, and evidence evaluation processing. These networks interacted with distinct cortical networks during encoding and retrieval, and differences in memory performance were associated with the spatiotemporal precision of these interactions. Longitudinally, instability of encoding-phase cerebellar–cerebral dynamics predicted reduced retrieval efficiency, particularly for old-item recognition. These findings extend models of memory ageing beyond cortico-hippocampal system and highlight dynamic cerebellar–cerebral coordination as a potential marker of memory resilience in later life.

## Methods

### Participants and study cohort

This study used neuroimaging and behavioural data from the Beijing Aging Brain Rejuvenation Initiative (BABRI), a longitudinal community-based cohort designed to identify early markers of cognitive impairment^40^. The initial cohort comprised 1,060 middle-aged and older adults aged 50 years or older. Participants were screened for neurological and psychiatric disorders, traumatic brain injury and major systemic conditions that could affect cognition. Dementia was excluded according to the Diagnostic and Statistical Manual of Mental Disorders, Fourth Edition, Revised (DSM-IV-R). Mild cognitive impairment was excluded according to Petersen’s diagnostic framework^41^. All participants included in the present analyses were cognitively normal, right-handed and had normal or corrected-to-normal vision.

The final cross-sectional sample included 821 participants aged 66.9 ± 7.1 years, including 570 women, with 11.5 ± 3.2 years of education. A longitudinal subset of 78 participants completed follow-up assessment after 4.0 ± 1.2 years. This subset had a baseline age of 66.0 ± 5.7 years, included 53 women and had 11.7 ± 2.9 years of education.

At baseline, participants were stratified into three groups according to quartiles of overall retrieval accuracy. The high- and low-performance groups included participants above or equal to the 75th percentile (= 0.89) and below or equal to the 25th percentile (= 0.75), respectively, the moderate-performance group included those within the interquartile range. Among participants with follow-up data, maintainers had stable or improved retrieval accuracy, whereas decliners had lower accuracy at follow-up.

### Episodic memory task

Before scanning, participants completed a 10–15 min practice session to ensure task comprehension. The main task included six encoding blocks and six retrieval blocks, each followed by fixation. Each task block contained six trials. The full fMRI task lasted 606 s, including instructions. During encoding, participants viewed 36 objects, including 16 natural and 20 man-made objects. During retrieval, they viewed 36 objects, including 18 studied items and 18 novel foils. The retrieval set was matched to the encoding set in category composition, both studied and novel items included 8 natural and 10 man-made objects. Stimuli were white line drawings presented on a black background using E-Prime 1.0. Visual stimuli were projected onto a head-coil-mounted screen, and responses were recorded via thumb presses. During encoding, participants responded with the left hand for natural objects and the right hand for man-made objects, during retrieval, they responded with the left hand for old items and the right hand for new items.

### MRI acquisition and preprocessing

MRI data were acquired on a Siemens TRIO 3T scanner at the Imaging Center for Brain Research, Beijing Normal University. Participants lay supine with head motion minimized using straps and foam pads. High-resolution T1-weighted structural images were acquired using a 3D magnetization-prepared rapid gradient-echo sequence for anatomical co-registration. Functional images were acquired using a T2-weighted echo-planar imaging sequence covering the whole brain. Scanning parameters can be found in Supplementary Text 1. Neuroimaging data were organized in BIDS format and preprocessed using DeepPrep, a GPU-accelerated pipeline optimized for large-scale datasets^42^. Preprocessing included slice-timing correction, distortion correction, coregistration, normalization, unwarping, confound estimation, segmentation and skull stripping. Normalization was performed to MNI152NLin6Asym space.

Nuisance regression was performed using Nilearn in Python and included 24 head-motion parameters (6 rigid-body motion estimates, their temporal derivatives, and quadratic terms of each) alongside white-matter, cerebrospinal-fluid, and global signal components (each expanded with temporal derivatives and quadratic terms), followed by linear detrending and z-score standardization. No spatial smoothing was applied at this stage, in order to preserve fine-grained cerebellar functional topography, as cerebellar folia and functional boundaries can be smaller than standard smoothing kernels^30^. The first 6 and last 3 TRs of each run were discarded to remove non-steady-state signal and end-of-run periods, respectively, leaving 294 volumes from the onset of the first encoding block to the end of the final retrieval fixation block.. Participants or runs were excluded for excessive head motion, defined as translation greater than 3 mm or rotation greater than 3 degrees, or incomplete cerebellar coverage below 90%.

### Task-evoked activation analysis

Participant-level GLMs were used to estimate task-evoked activation maps from the experimental design, followed by group-level models to assess population-level effects. Three classes of contrasts were examined: (1) memory-phase contrasts (encoding versus fixation and retrieval versus fixation), (2) retrieval stimulus-type contrasts (old items versus fixation, new items versus fixation, and old versus new items), and (3) a motor-response contrast (left-hand versus right-hand responses).

At the first level, we modelled the haemodynamic response using the Glover canonical double-gamma response function, accounted for temporal autocorrelation with an AR(1) noise model, and modelled low-frequency signal drifts using a cosine-basis set. Group-level models included age, gender and years of education as covariates. Voxel-wise statistical maps were FDR corrected, with significance defined as FDR-adjusted P < 0.05. For network-level analyses involving component-based measures, we applied the more stringent Bonferroni correction to control for multiple comparisons.

### Cerebellar functional parcellation with sparse dictionary learning

To identify memory-relevant cerebellar organization without imposing predefined anatomical or functional boundaries, we applied sparse dictionary learning to task-activated cerebellar voxels^43^. This approach allows spatially overlapping components and is well suited to the cerebellum, where functional fields can be compact and partially interdigitated. The analysis was restricted to the intersection of the group-level task-responsive mask and an anatomical cerebellar mask provided by the Spatially Unbiased Infratentorial Template (SUIT), minimizing contamination from adjacent occipital cortex^44^.

We optimized the number of components and the sparsity regularization parameter using grid search, with both parameters ranging from 10 to 70 in increments of 5. The model used a smoothing kernel of 4 *mm*, 50 training epochs, and a batch size of 20. We applied the Kneedle algorithm sequentially: first, we identified the optimal component number from the elbow of the explained-variance curve using a concave, increasing function, then, we identified the optimal sparsity parameter using a decreasing curve. This procedure balanced model complexity against the variance explained by the decomposition. After excluding three artifactual brainstem components, 47 cerebellar components were retained for subsequent analyses (Extended Data Fig. 1). The large sample size was expected to support stable decomposition, consistent with prior dimensionality-reduction-based fMRI analyses in samples exceeding 700 participants^35^.

### Selection of memory-relevant cerebellar networks

We identified memory-relevant cerebellar networks using a three-step procedure. First, for each cerebellar network, we quantified its spatial overlap with the activation map for the left-versus right-hand response thresholded at FDR-adjusted *P* < 0.05. We defined the overlap ratio as the proportion of voxels within the network that were also included in the thresholded activation map. Networks with an overlap ratio greater than 50% were classified as predominantly sensorimotor and excluded. Second, we excluded networks that showed significant activation in the hand-response contrast. Finally, we tested the remaining networks for associations between contrast-specific network activation and behavioural measures, including encoding and retrieval accuracy, response time, discrimination sensitivity and response criterion. These associations were assessed using GLMs with age, sex and education as covariates. Networks showing a significant association with at least one behavioural measure (*P* <0.05 after Bonferroni correction) were retained as memory-relevant cerebellar networks for subsequent analyses.

### HRF-informed temporal brain–behaviour modulation

To quantify trial-level associations between cerebellar activity and behaviour, we fitted an HRF-informed first-level GLMs separately for each participant, cerebellar network and task phase. Encoding and retrieval were modelled independently using three regressors: a task regressor, a trial-level accuracy modulator and a trial-level response-time modulator. Encoding accuracy was defined by subsequent memory performance, whereas retrieval accuracy reflected correct old/new recognition. Response time series were z-scored within participant across both phases, and the accuracy and response-time modulators were mean-centred within participant and phase before being entered simultaneously into the model. Trial onsets were represented at TR resolution, and each trial was modelled as a 4-s boxcar event. All regressors were convolved with an SPM-like canonical double-gamma HRF. Fixation periods were not modelled as behavioural events and contributed to baseline and drift estimation.

For each participant, network and phase, we fitted the following model:

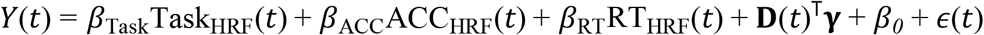

where *Y*(*t*) denotes the cerebellar network time series; *β*_Task_, *β*_ACC_, and *β*_RT_ denote the regression coefficients for the task, accuracy, and response-time regressors, respectively; Task_HRF_(*t*), ACC_HRF_(*t*), and RT_HRF_(*t*) denote the corresponding HRF-convolved regressors; **D**(*t*) denotes the vector of discrete cosine drift regressors; **γ** denotes the corresponding vector of regression coefficients; *β*_*0*_ denotes the intercept; and *ϵ*(*t*) denotes the residual error. Low-frequency drifts were modelled using a discrete cosine basis with a 128-s high-pass cutoff.

To account for temporal autocorrelation, we estimated and constrained an AR(1) coefficient from the residuals of an initial ordinary least-squares fit, used it to prewhiten the data and design matrix, and refitted the model. For each participant, network and phase, we extracted beta estimates, t-statistics and P-values for the task, accuracy and response-time regressors. Group-level effects were assessed using one-sample t-tests against zero within each performance group, with FDR correction applied across tests.

### Static and dynamic functional connectivity

Functional connectivity was quantified between memory-relevant cerebellar networks and cortical systems defined by the Schaefer-400 atlas^45^ and Yeo 17-network parcellation^46^. Static functional connectivity was estimated separately for encoding and retrieval using Ledoit–Wolf shrinkage correlations across all volumes within each phase.

Dynamic functional connectivity was estimated using sliding-window correlations with Ledoit–Wolf shrinkage. The window length was 24 s, corresponding to one task block, and the step size was 1 TR. Although resting-state dynamic connectivity studies often use longer windows to reduce low-frequency noise^47^, the present task-based design required a window length matched to the experimental structure. Prior work shows that windows of approximately 22.5 s can capture ongoing task-related cognitive states^48^. The Schaefer-400 atlas was used to provide sufficient spatial granularity for short-window dynamic functional connectivity analyses^48^.

### Identification of recurrent dynamic connectivity states

To identify recurrent cerebellar–cerebral connectivity configurations, we applied K-means clustering to sliding-window dynamic connectivity matrices^49^. For each window, connectivity values between the 3 cerebellar networks and 400 cortical parcels were concatenated into a 1200-dimensional vector. Before clustering, the 1% of samples with the largest absolute connectivity values were removed as extreme outliers, and the remaining data were standardized across features using z scores. Candidate solutions from k = 2 to k = 5 were evaluated using the silhouette coefficient, with 20 random initializations for each solution^50,51^.

### Dynamic time warping analysis

Dynamic time warping (DTW) was used to compare dynamic connectivity trajectories across performance groups and across longitudinal time points^52^. This method quantifies sequence similarity by identifying an optimal nonlinear temporal alignment and is less sensitive than point-by-point correlation to phase offsets and temporal lags^53^. Larger DTW distance indicated greater divergence. For each memory-relevant cerebellar network and each cortical parcel, DTW distances were computed between participants’ dynamic functional connectivity trajectories within each task phase. Dynamic connectivity trajectories were derived from sliding-window connectivity estimates using a 12-TR window, corresponding to 24 s, and a step size of 1 TR. This window length matched the duration of one task block and was used to preserve temporal correspondence with the task structure.

For cross-sectional performance-group analyses, pairwise DTW distances were computed between participants from each pair of performance groups, including high versus low, high versus moderate and moderate versus low. The median pairwise DTW distance was used as the observed summary measure of between-group trajectory divergence for each cerebellar network–cortical parcel pair. Larger median DTW values indicated greater divergence in the temporal shape or alignment of cerebellar–cerebral coupling trajectories between groups. For longitudinal analyses, DTW distances were computed between baseline and follow-up trajectories within participants, with larger values indicating lower longitudinal stability of dynamic coupling.

Before DTW analysis, we regressed age, sex and education from the dynamic connectivity time series. For each task phase, observed median between-group DTW distances were standardized against the empirical distribution across all tested cerebellar network–cortical parcel pairs. Connections with an absolute standardized score of *P* > 1.96 were retained, corresponding to a two-sided threshold of *Z* < 0.05. For each retained connection, a non-parametric permutation test with 5,000 iterations was then performed by randomly shuffling group labels while preserving group sizes. In each iteration, the median between-group DTW distance was recomputed to generate a null distribution. The two-sided permutation *P* value was calculated as the proportion of permuted distances at least as extreme as the observed distance. A connection was considered to show a permutation-screened trajectory difference if it met both criteria: |*Z*| > 1.96 and *P* < 0.05.

### Statistical analysis

fMRI task performance was summarized by accuracy and response time during encoding and retrieval, including separate measures for old and new items during retrieval. Baseline group differences were assessed using one-way analyses of variance (ANOVAs) for age and years of education and a chi-square test for gender. Group differences in task performance were tested using analyses of covariance (ANCOVAs) adjusted for age, gender and education. Paired t-tests assessed old–new differences during retrieval and longitudinal changes in all behavioural measures between baseline and follow-up. Discrimination sensitivity and response criterion were calculated using signal-detection theory.

The behavioural relevance of cerebellar network activation was assessed using GLMs relating contrast-specific network activation to task performance, with Bonferroni correction for multiple comparisons. Group differences in static functional connectivity and state occupancy were tested using ANCOVAs, with age, sex and education included as covariates, followed by FDR correction across cerebral parcels.

Longitudinal changes were defined as follow-up minus baseline. Differences between baseline and follow-up in static functional connectivity were assessed using paired t-tests, followed by FDR correction. Associations between longitudinal changes in functional connectivity and behavioural performance were examined using GLMs relating functional connectivity changes to behavioural changes. Dynamic stability was quantified using DTW distance between baseline and follow-up connectivity trajectories, with larger distances indicating greater longitudinal instability.

## Data availability

The raw behavioural and neuroimaging data used in this study are part of the Beijing Aging Brain Rejuvenation Initiative (BABRI) cohort^40^. Data are available upon reasonable request to the corresponding author, subject to the data sharing regulations of the BABRI project. All derived data supporting the findings of this study, including unthresholded and thresholded group-level statistical brain maps, the functional parcellation of cerebellar networks and statistical result tables, are publicly available in the GitHub repository (https://github.com/Tengfei-dare/cerebellum-networks-memory). The whole-brain mask is retrieved via TemplateFlow (https://www.templateflow.org/). The cerebellar mask and atlas are provided by SUIT (https://www.diedrichsenlab.org/imaging/atlasPackage.htm). The cortical Schaefer atlas used in this study was retrieved via the nilearn library (https://nilearn.github.io/stable/modules/datasets.html#deterministic-atlases).

## Code availability

All custom analytical scripts developed for this study, including scripts for task-related activation analysis, sparse dictionary learning, dynamic functional connectivity estimation, DTW distance calculation and statistical visualization, are available at GitHub (https://github.com/Tengfei-dare/cerebellum-networks-memory). The analyses were primarily executed using Python v3.12.3, utilizing Nilearn v0.12.0, SciPy v1.16.1, DTAIDistance v2.3.13. Detailed documentation regarding the DeepPrep(v25.1.0) neuroimaging preprocessing pipeline can be accessed at (https://deepprep.readthedocs.io/en/latest/).

## Acknowledgements

This work was supported by Brain Science and Brain - like Intelligence Technology - National Science and Technology Major Project(Grant No. 2022ZD0211600), Noncommunicable Chronic Diseases-National Science and Technology Major Project (Grant No. 2025ZD0546300), and Joint Innovation Team for Clinical & Basic Research,Shandong First Medical University (No. CX202408), the Major Project of National Social Science Foundation (24&5ZD252), and Interdisciplinary Research Foundation for Doctoral Candidates of Beijing Normal University (Grant BNUXKJC2410).

We thank all the volunteers, the clinical and neuroimaging staff for their participation and support.

## Author contributions

Tengfei Han, Zaizhu Han, Yaojing Chen, and Zhanjun Zhang conceptualized and designed this study. Tengfei Han, Mingxi Dang, Wenhao Bai, Ziyun Li, Chi Zhang, Yaojing Chen, Zhanjun Zhang collected the behavioural and imaging data. Tengfei Han and Ziyun Li preprocessed the data. Tengfei Han performed the data analysis and interpretation under the supervision of Zaizhu Han, Yaojing Chen, and Zhanjun Zhang. Tengfei Han drafted the first version of the manuscript, and Ziyun Li, Chi Zhang, Yaojing Chen, Zaizhu Han provided critical revisions.

---

**Extended Data Fig. 1.**
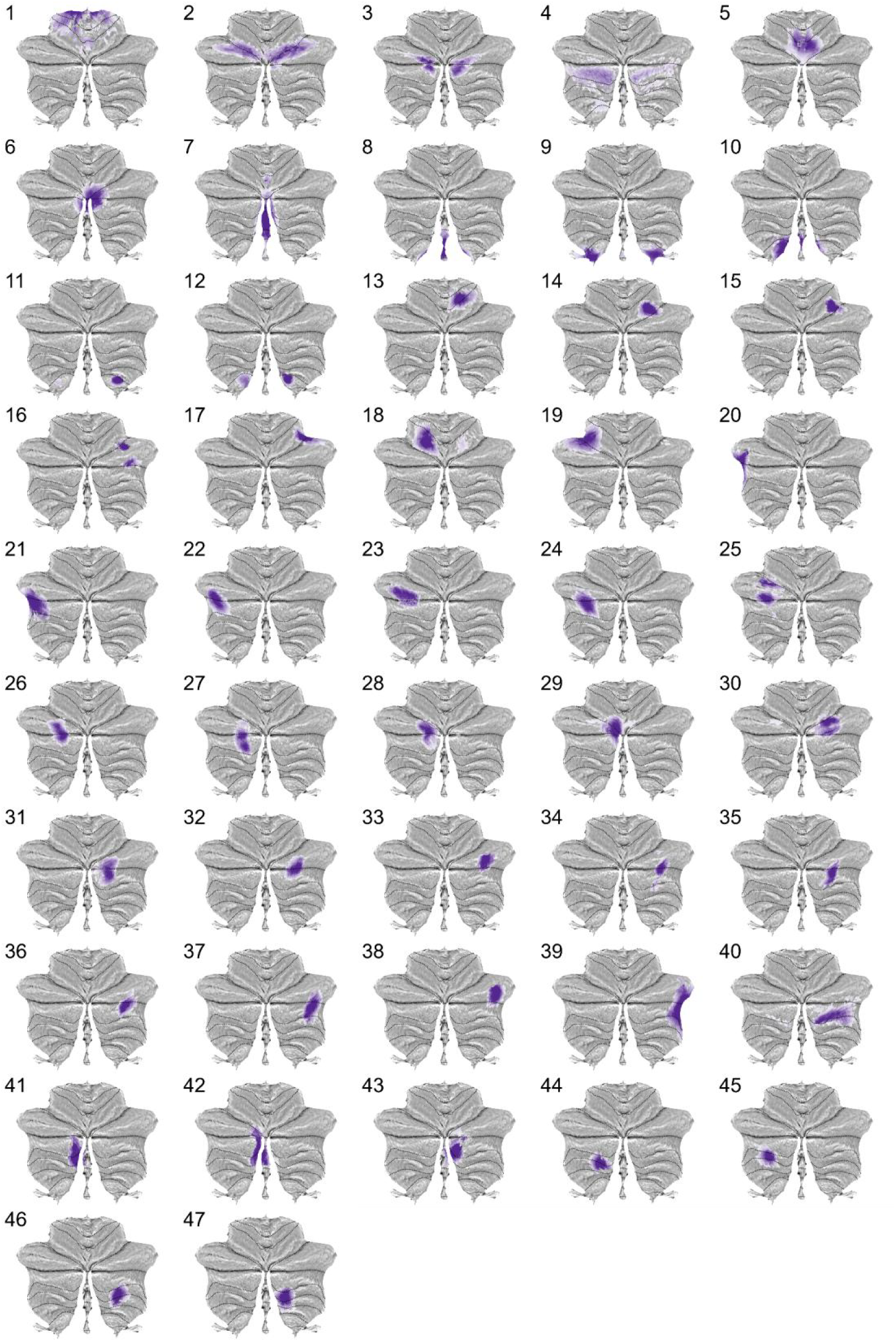
Task-responsive cerebellar functional networks derived by sparse dictionary learning. Approximately 40% networks were located within Crus I and Crus II and were characterized mainly by multiple spatially distinct unilateral representations (Nets 20–40). Bilateral components followed an anterior-to-posterior gradient, localizing to the lobule VI/Crus I boundary (Net 2), medial Crus I/II (Net 3), the Crus II/lobule VIIb interface (Net 4) and lobule IX (Nets 9–12).

**Extended Data Table 1.** Demographic characteristics and episodic memory task performance at baseline. Values are presented as mean (SD) unless otherwise indicated. Participants were stratified into High, Moderate, and Low groups based on their baseline memory retrieval performance, according to quartile. Group differences in demographic characteristics were assessed using one-way analysis of variance (ANOVA) for age and education, and a two-sided Pearson’s *χ*^*2*^ test for sex. Differences in encoding and retrieval task performance across groups were evaluated using analysis of covariance (ANCOVA), adjusting for age, gender, and years of education as covariates. ACC, accuracy; RT, reaction time.

| | All | High | Moderate | Low | $F/\chi^2$ | $p$ |
| --- | --- | --- | --- | --- | --- | --- |
| n | 821 | 299 | 271 | 251 |  |  |
| Age, years | 66.9 (7.1) | 66.2 (6.5) | 66.0 (6.7) | 68.8 (7.7) | 13.292 | <0.001 |
| Female, n (%) | 570 (69.4) | 213 (71.2) | 193 (71.2) | 164 (65.3) | 3.384 | 0.184 |
| Education, years | 11.5 (3.2) | 11.9 (2.9) | 11.6 (2.8) | 11.0 (3.6) | 4.993 | 0.007 |
| **Encoding** |  |  |  |  |  |  |
| Overall ACC | 0.94 (0.11) | 0.97 (0.07) | 0.95 (0.08) | 0.90 (0.16) | 23.327 | <0.001 |
| Mean RT, ms | 1272 (358) | 1215 (279) | 1246 (290) | 1395 (485) | 15.540 | <0.001 |
| **Retrieval** |  |  |  |  |  |  |
| Old item ACC | 0.70 (0.18) | 0.85 (0.07) | 0.70 (0.09) | 0.51 (0.17) | 563.382 | <0.001 |
| New item ACC | 0.89 (0.14) | 0.94 (0.06) | 0.91 (0.08) | 0.79 (0.22) | 83.453 | <0.001 |
| Old item RT, ms | 1561 (357) | 1418 (248) | 1557 (265) | 1765 (455) | 62.498 | <0.001 |
| New item RT, ms | 1467 (388) | 1374 (251) | 1419 (273) | 1671 (553) | 33.084 | <0.001 |
| Overall ACC | 0.80 (0.12) | 0.90 (0.04) | 0.81 (0.02) | 0.65 (0.13) | 649.763 | <0.001 |
| Mean RT, ms | 1514 (341) | 1396 (220) | 1486 (232) | 1717 (466) | 54.534 | <0.001 |
| $d'$ | 1.85 (0.69) | 2.44 (0.40) | 1.82 (0.31) | 1.13 (0.60) | 520.369 | <0.001 |
| Criterion | 0.45 (0.36) | 0.30 (0.25) | 0.46 (0.31) | 0.60 (0.46) | 56.847 | <0.001 |

