## Supplemental figures, tables, texts for "Spatiotemporal coordination of specialized cerebellar networks supports episodic memory in older adults"

**Supplementary File**

### **Supplementary Text 1. MRI acquisition parameters.**

MRI data were acquired on a Siemens TRIO 3T scanner (Berlin, Germany) at the Imaging Center for Brain Research, Beijing Normal University. High-resolution T1-weighted structural images were acquired using a 3D magnetization-prepared rapid gradient-echo (MPRAGE) sequence (176 slices, TR = 1900 ms, TE = 3.44 ms, slice thickness = 1 mm, FA = 9°, FOV = 256×256 mm², acquisition matrix = 256×256) for anatomical co-registration. Functional images employed a T2-weighted echo-planar imaging (EPI) sequence (33 slices, repetition time (TR) = 2000 ms, echo time (TE) = 30 ms, slice thickness = 3.5 mm, flip angle (FA) = 90°, field of view (FOV) = 200×200 mm², acquisition matrix = 64×64) covering the whole brain.

### **Supplementary Fig. 1. Task-evoked activation profiles of cerebellar functional networks across task conditions.**

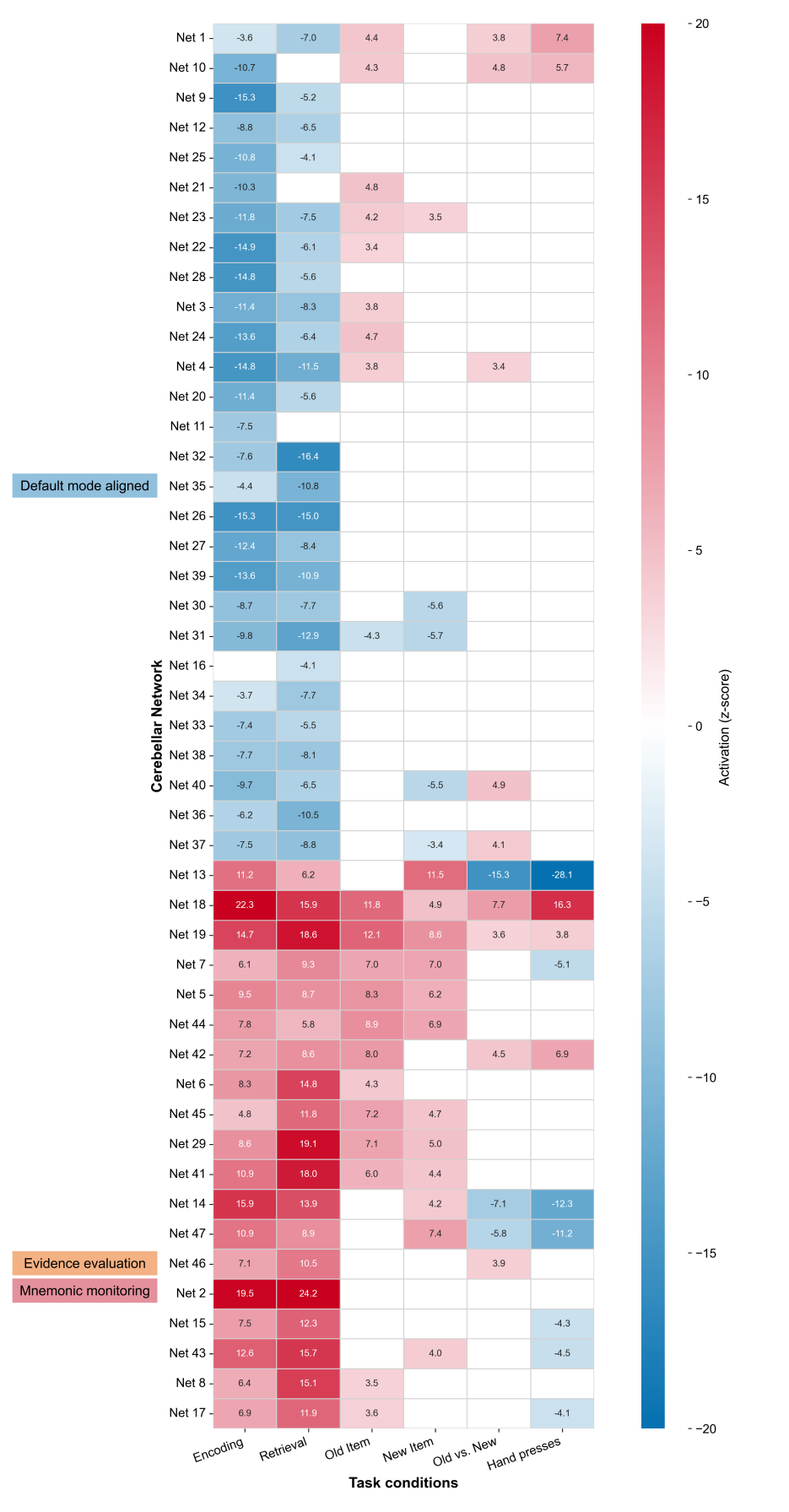

To facilitate visualization of task-related functional profiles, networks were ordered using hierarchical clustering based on their activation patterns across the six task conditions.

### **Supplementary Fig. 2. Hyperparameter sensitivity of the Net 2 cerebellar parcellation.**

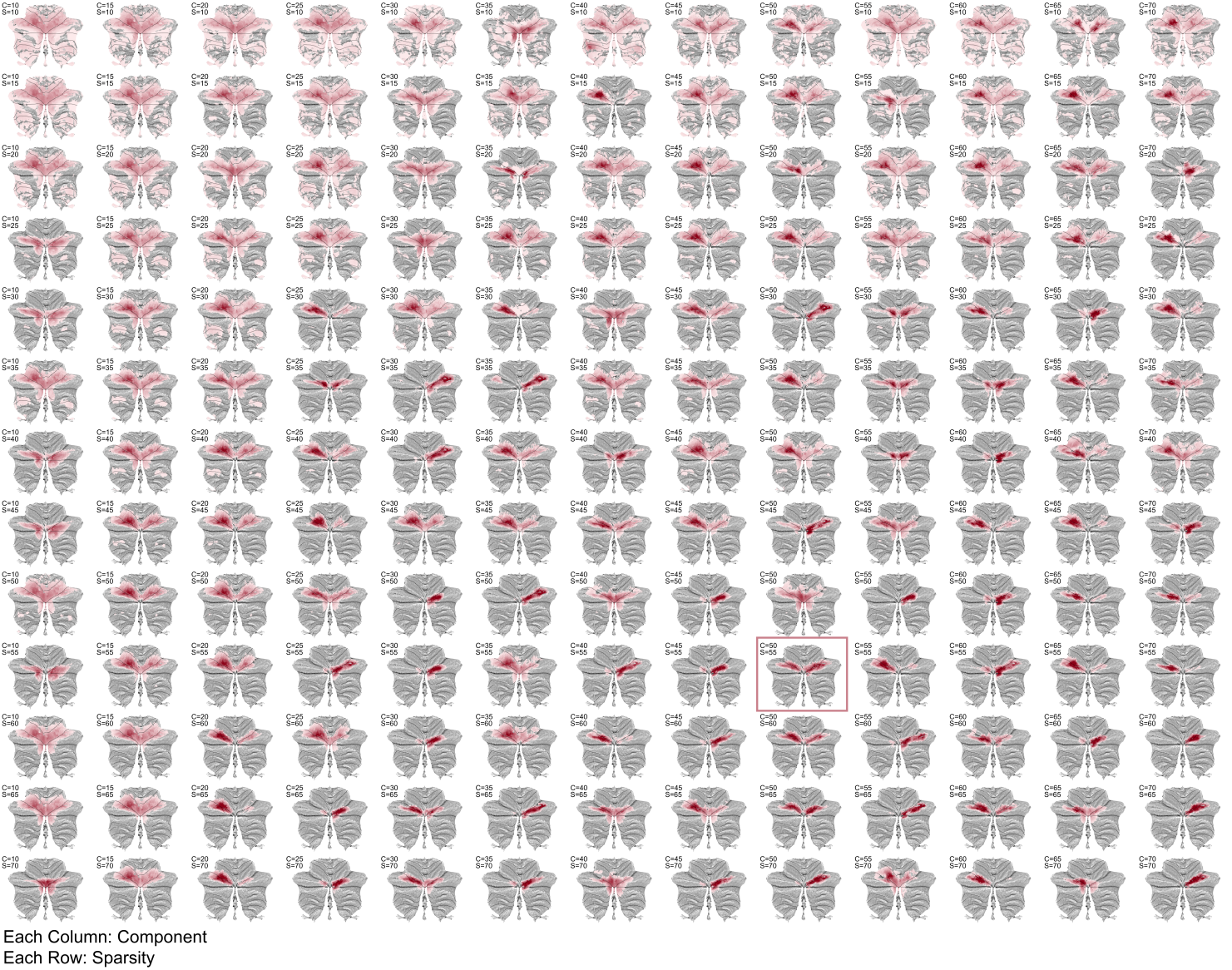

Sparse dictionary learning was repeated across a grid of hyperparameters to assess the robustness of Net 2. For each combination of the number of components C and the sparsity parameter S, the component most similar to the reference Net 2 was identified using the Dice coefficient. Columns indicate increasing numbers of components from 10 to 70 in increments of 5, and rows indicate increasing sparsity values from 10 to 70 in increments of 5.

### **Supplementary Fig. 3. Hyperparameter sensitivity of the Net 35 cerebellar parcellation.**

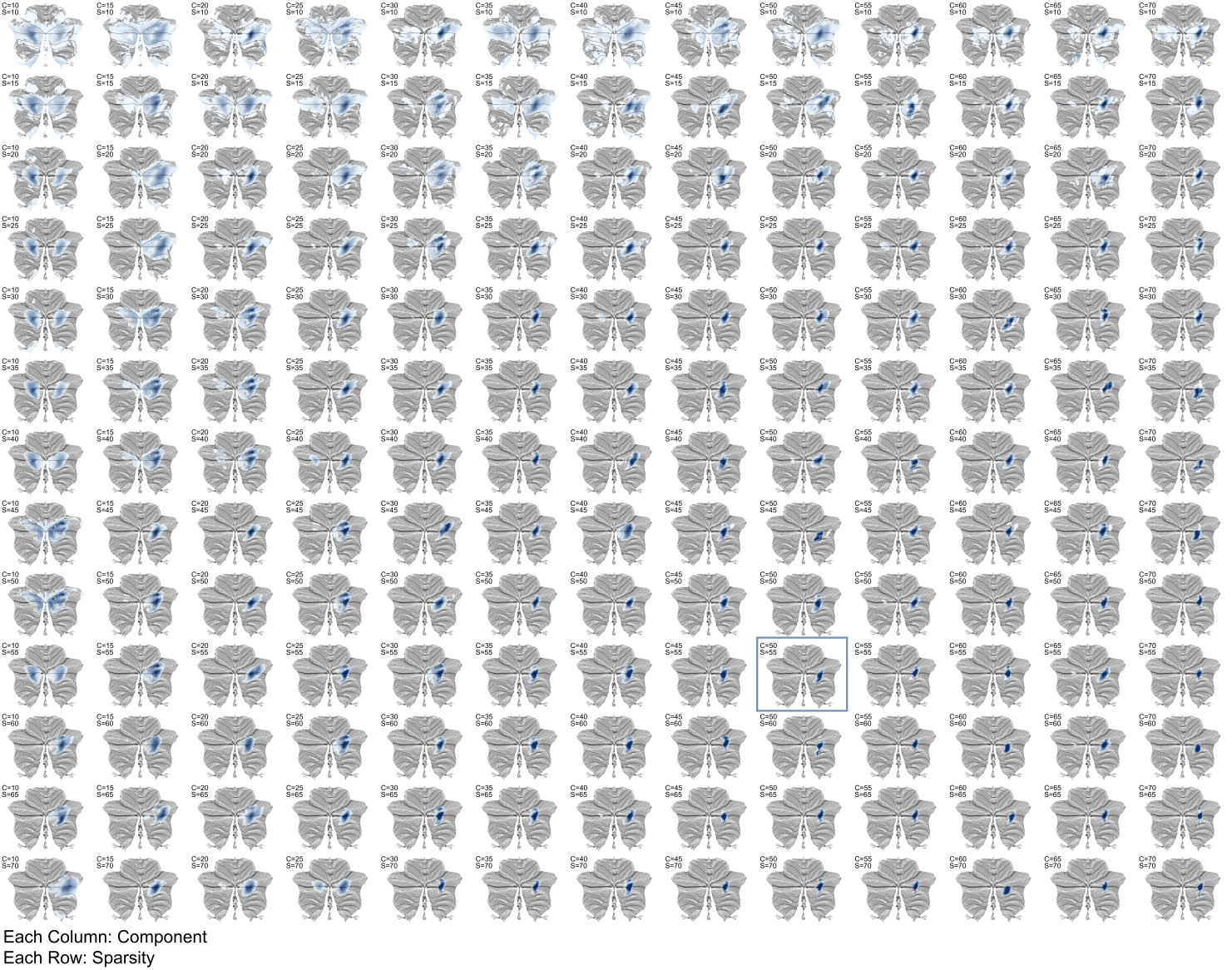

Sparse dictionary learning was repeated across a grid of hyperparameters to assess the robustness of Net 35. For each combination of the number of components C and the sparsity parameter S, the component most similar to the reference Net 35 was identified using the Dice coefficient. Columns indicate increasing numbers of components from 10 to 70 in increments of 5, and rows indicate increasing sparsity values from 10 to 70 in increments of 5.

### **Supplementary Fig. 4. Hyperparameter sensitivity of the Net 46 cerebellar parcellation.**

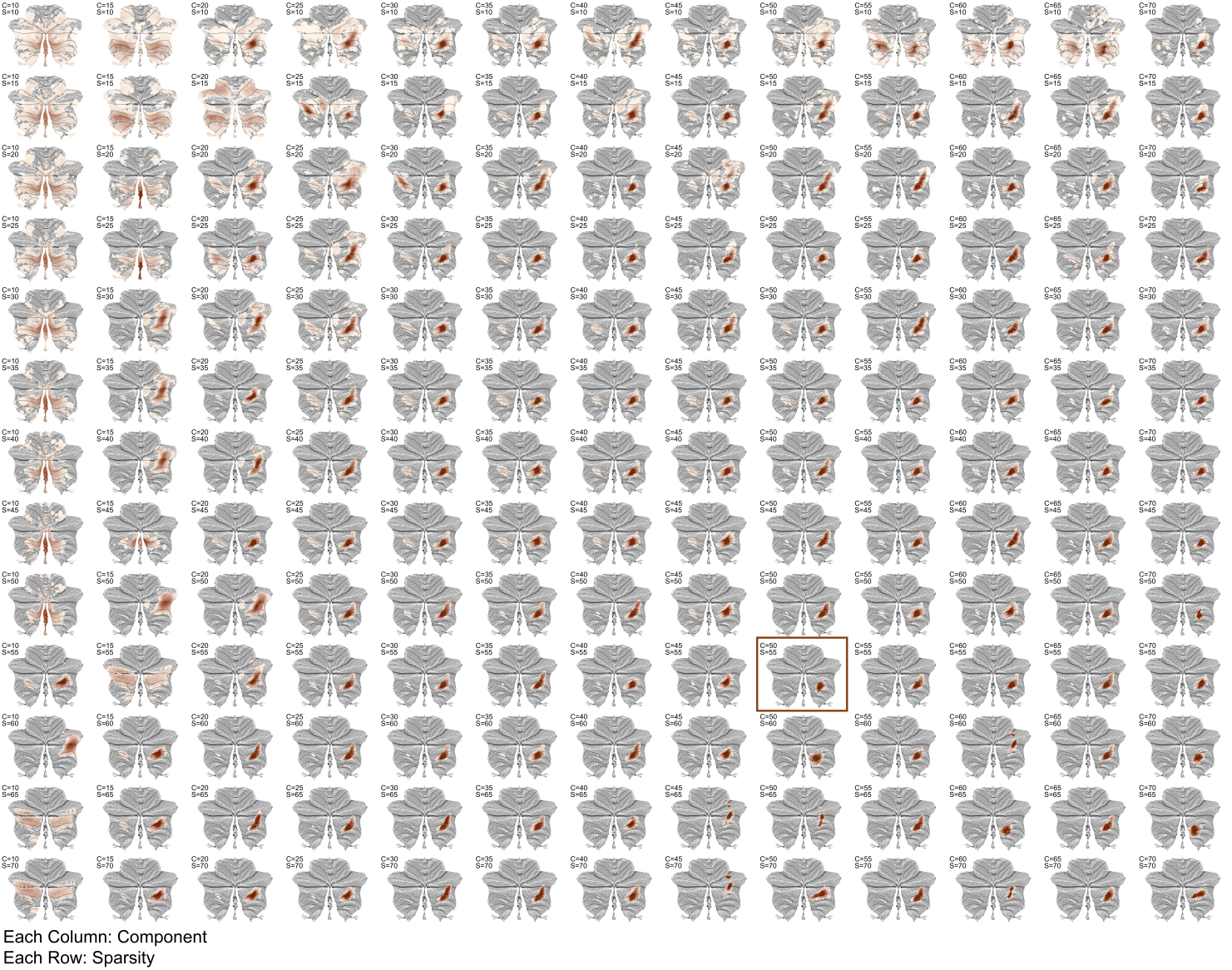

Sparse dictionary learning was repeated across a grid of hyperparameters to assess the robustness of Net 46. For each combination of the number of components C and the sparsity parameter S, the component most similar to the reference Net 46 was identified using the Dice coefficient. Columns indicate increasing numbers of components from 10 to 70 in increments of 5, and rows indicate increasing sparsity values from 10 to 70 in increments of 5.

### **Supplementary Fig. 5. Silhouette coefficients in k-means clustering of sliding-window cerebellar–cerebral dynamic connectivity.**

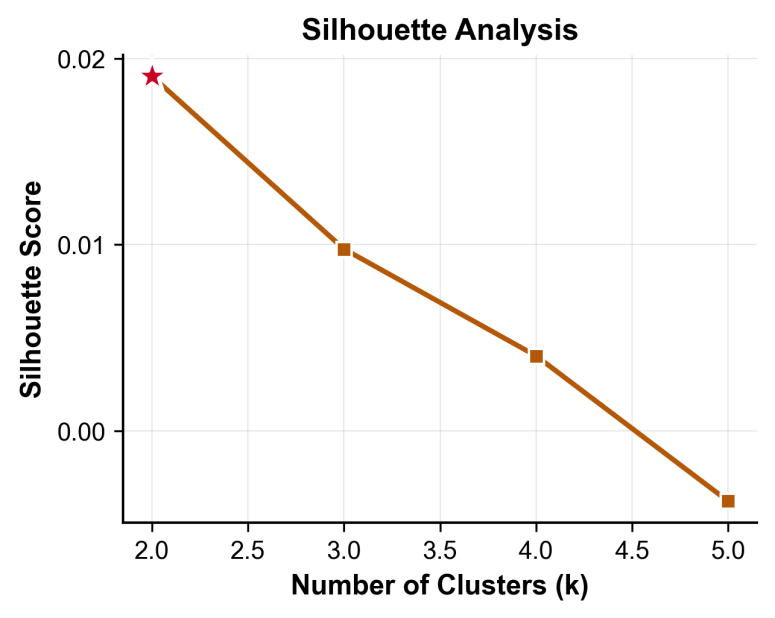

### **Supplementary Fig. 6. Correspondence between memory-related cerebellar networks and established functional and gradient-based cerebellar atlases.**

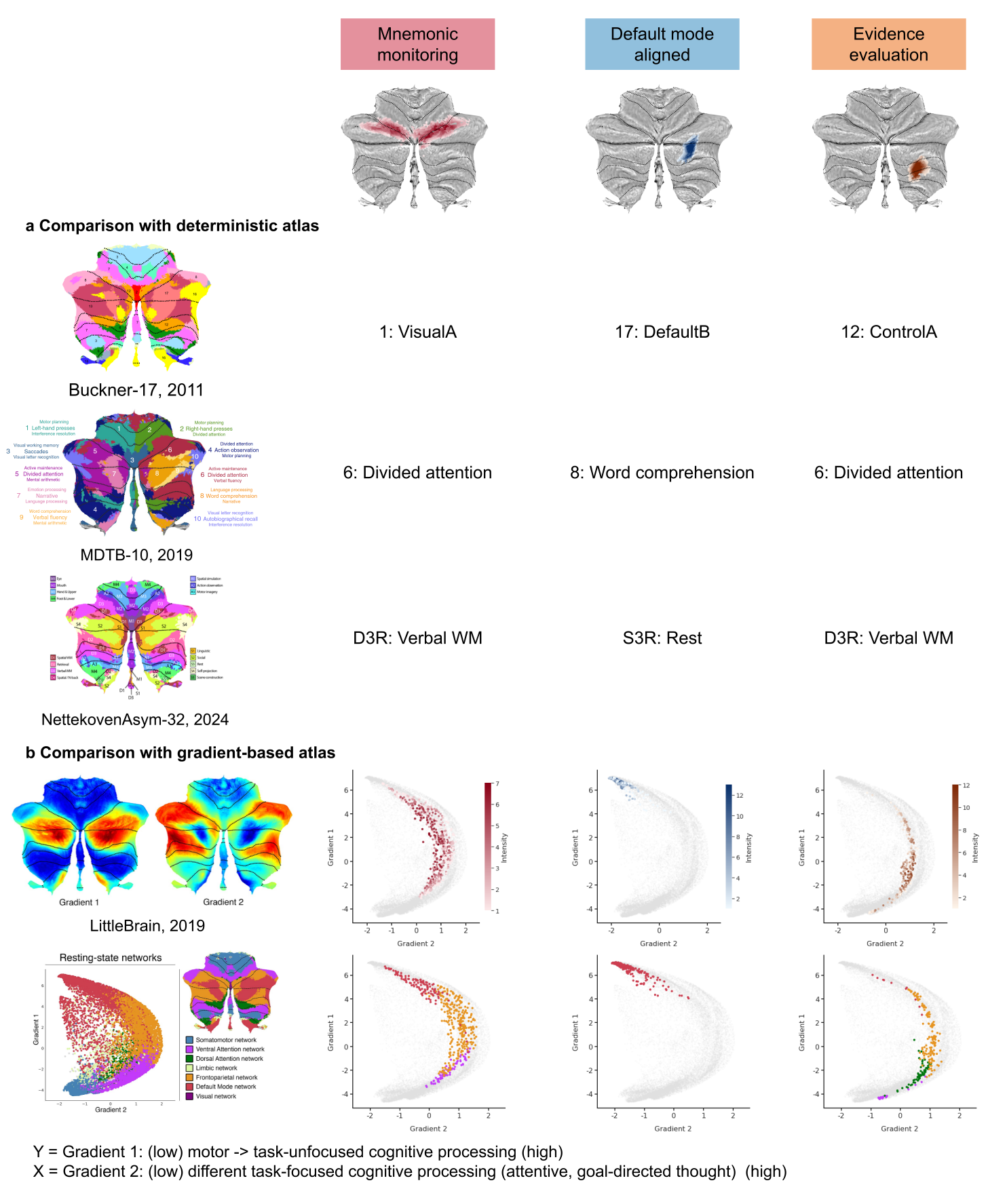

The spatial maps of the identified cerebellar components were compared with those from published cerebellar atlases using Dice coefficients.

### **Supplementary Table 1. Demographic characteristics and episodic memory task performance in the longitudinal follow-up sample.**

|  | Baseline | Follow-up | *t* | *p* |
| --- | --- | --- | --- | --- |
| n | 78 | 78 | - | - |
| Age, years | 66.0 (5.7) | 69.9 (5.8) | -28.157 | <0.001 |
| Female, n (%) | 53 (67.9) | - | - | - |
| Education, years | 11.7 (2.9) | - | - | - |
| **Encoding** |  |  |  |  |
| Overall ACC | 0.96 (0.10) | 0.97 (0.05) | -0.534 | 0.595 |
| Mean RT, ms | 1179 (318) | 1178 (269) | 0.034 | 0.973 |
| **Retrieval** |  |  |  |  |
| Old item ACC | 0.73 (0.15) | 0.72 (0.16) | 0.434 | 0.665 |
| New item ACC | 0.91 (0.08) | 0.93 (0.07) | -1.601 | 0.113 |
| Old item RT, ms | 1451 (287) | 1470 (306) | -0.559 | 0.578 |
| New item RT, ms | 1319 (216) | 1341 (250) | -0.747 | 0.457 |
| Overall ACC | 0.82 (0.08) | 0.83 (0.08) | -0.418 | 0.677 |
| Mean RT, ms | 1382 (227) | 1405 (248) | -0.820 | 0.415 |
| d’ | 1.87 (0.55) | 2.03 (0.51) | -2.140 | 0.036 |
| Criterion | 0.41 (0.32) | 0.45 (0.34) | -0.885 | 0.379 |

Values are presented as mean (SD) unless otherwise indicated. ACC, accuracy; RT, reaction time. t values and p values are derived from paired-sample t-tests.

### **Supplementary Table 2 Demographic characteristics and episodic memory task performance in the longitudinal maintainers sample.**

|  | Baseline | Follow-up | *t* | *p* |
| --- | --- | --- | --- | --- |
| n | 48 | 48 | - | - |
| Age, years | 66.0 (5.7) | 69.9 (5.8) | -20.941 | <0.001 |
| Female, n (%) | 31 (64.6) | - | - | - |
| Education, years | 11.7 (2.8) | - | - | - |
| **Encoding** |  |  |  |  |
| Overall ACC | 0.95 (0.11) | 0.96 (0.06) | -0.238 | 0.813 |
| Mean RT, ms | 1187 (297) | 1199 (303) | -0.282 | 0.779 |
| **Retrieval** |  |  |  |  |
| Old item ACC | 0.71 (0.16) | 0.77 (0.14) | -4.530 | <0.001 |
| New item ACC | 0.89 (0.09) | 0.94 (0.07) | -3.633 | <0.001 |
| Old item RT, ms | 1464 (300) | 1418 (290) | 1.271 | 0.210 |
| New item RT, ms | 1312 (220) | 1318 (234) | -0.175 | 0.862 |
| Overall ACC | 0.80 (0.07) | 0.86 (0.06) | -8.028 | <0.001 |
| Mean RT, ms | 1385 (236) | 1368 (222) | 0.564 | 0.575 |
| d’ | 1.71 (0.52) | 2.21 (0.42) | -7.764 | <0.001 |
| Criterion | 0.41 (0.36) | 0.39 (0.34) | 0.420 | 0.676 |

Values are presented as mean (SD) unless otherwise indicated. ACC, accuracy; RT, reaction time. t values and p values are derived from paired-sample t-tests.

### **Supplementary Table 3. Demographic characteristics and episodic memory task performance in the longitudinal decliners sample.**

|  | Baseline | Follow-up | *t* | *p* |
| --- | --- | --- | --- | --- |
| n | 30 | 30 | - | - |
| Age, years | 65.9 (5.8) | 70.0 (5.8) | -18.956 | <0.001 |
| Female, n (%) | 22 (73.3) | - | - | - |
| Education, years | 11.7 (3.1) | - | - | - |
| **Encoding** |  |  |  |  |
| Overall ACC | 0.97 (0.08) | 0.98 (0.03) | -0.652 | 0.519 |
| Mean RT, ms | 1167 (353) | 1145 (202) | 0.408 | 0.687 |
| **Retrieval** |  |  |  |  |
| Old item ACC | 0.77 (0.13) | 0.64 (0.16) | 7.278 | <0.001 |
| New item ACC | 0.94 (0.05) | 0.91 (0.08) | 1.811 | 0.081 |
| Old item RT, ms | 1429 (269) | 1553 (318) | -1.959 | 0.060 |
| New item RT, ms | 1330 (213) | 1377 (273) | -0.912 | 0.369 |
| Overall ACC | 0.86 (0.07) | 0.78 (0.08) | 9.186 | <0.001 |
| Mean RT, ms | 1378 (217) | 1463 (278) | -1.709 | 0.098 |
| d’ | 2.13 (0.51) | 1.73 (0.51) | 4.906 | <0.001 |
| Criterion | 0.42 (0.24) | 0.54 (0.33) | -2.084 | 0.046 |

Values are presented as mean (SD) unless otherwise indicated. ACC, accuracy; RT, reaction time. t values and p values are derived from paired-sample t-tests.

### **Supplementary Table 4. Anatomical localization and sensorimotor overlap of 47 cerebellar functional networks derived using sparse dictionary learning.**

| Network ID | Anatomical Location  (SUIT Atlas) | Hemisphere | Volume  (Voxels) | Included in Motor Activation Map (%) | Putative Category |
| --- | --- | --- | --- | --- | --- |
| Net 1 | I-V | Bilateral | 1118 | 38.55 | Cognitive |
| Net 2 | VI-CrusI | Bilateral | 926 | 11.45 | Cognitive |
| Net 3 | CrusI-CrusII | Bilateral | 697 | 0.57 | Cognitive |
| Net 4 | CrusII-VIIb | Bilateral | 1614 | 8.61 | Cognitive |
| Net 5 | V-VI | Bilateral | 739 | 55.35 | Motor |
| Net 6 | Vermal VI-CrusII | Bilateral | 487 | 3.29 | Cognitive |
| Net 7 | Vermal CrusI-VIIIb | Bilateral | 356 | 42.42 | Cognitive |
| Net 8 | Vermal VIIIb-IX | Bilateral | 358 | 10.89 | Cognitive |
| Net 9 | IX | Bilateral | 562 | 4.27 | Cognitive |
| Net 10 | Vermal IX | Bilateral | 334 | 26.05 | Cognitive |
| Net 11 | IX | Bilateral | 251 | 4.78 | Cognitive |
| Net 12 | IX | Bilateral | 320 | 20.00 | Cognitive |
| Net 13 | V-VI | Right | 372 | 99.19 | Motor |
| Net 14 | VI | Right | 235 | 58.72 | Motor |
| Net 15 | VI | Right | 200 | 28.00 | Cognitive |
| Net 16 | CrusI | Right | 281 | 2.14 | Cognitive |
| Net 17 | VI-CrusI | Right | 208 | 23.56 | Cognitive |
| Net 18 | VI | Left | 549 | 72.31 | Motor |
| Net 19 | VI-CrusI | Left | 631 | 19.65 | Cognitive |
| Net 20 | CrusI | Left | 309 | 0.65 | Cognitive |
| Net 21 | CrusI-CrusII | Left | 364 | 1.10 | Cognitive |
| Net 22 | CrusI-CrusII | Left | 417 | 0.96 | Cognitive |
| Net 23 | CrusI-CrusII | Left | 427 | 1.64 | Cognitive |
| Net 24 | CrusI-CrusII | Left | 480 | 1.25 | Cognitive |
| Net 25 | CrusI-CrusII | Left | 478 | 0.21 | Cognitive |
| Net 26 | CrusI-CrusII | Left | 524 | 1.15 | Cognitive |
| Net 27 | CrusI-CrusII | Left | 491 | 1.43 | Cognitive |
| Net 28 | CrusI-CrusII | Left | 524 | 0.57 | Cognitive |
| Net 29 | V-CrusI | Left | 509 | 5.89 | Cognitive |
| Net 30 | CrusI | Right | 474 | 1.69 | Cognitive |
| Net 31 | CrusI-CrusII | Right | 480 | 2.08 | Cognitive |
| Net 32 | CrusI-CrusII | Right | 391 | 1.02 | Cognitive |
| Net 33 | CrusI | Right | 220 | 6.36 | Cognitive |
| Net 34 | CrusI-CrusII | Right | 285 | 2.46 | Cognitive |
| Net 35 | CrusI-CrusII | Right | 317 | 0.63 | Cognitive |
| Net 36 | CrusI-CrusII | Right | 304 | 3.29 | Cognitive |
| Net 37 | CrusI-CrusII | Right | 328 | 4.27 | Cognitive |
| Net 38 | CrusI | Right | 236 | 2.54 | Cognitive |
| Net 39 | CrusI-CrusII-VIIb | Right | 408 | 1.47 | Cognitive |
| Net 40 | CrusII-VIIb | Right | 632 | 5.70 | Cognitive |
| Net 41 | CrusII-VIIb | Left | 269 | 13.38 | Cognitive |
| Net 42 | Vermal CrusII-VIIb | Left | 273 | 39.19 | Cognitive |
| Net 43 | Vermal CrusII-VIIb | Right | 350 | 28.86 | Cognitive |
| Net 44 | VIIb-VIIIa | Left | 253 | 15.42 | Cognitive |
| Net 45 | VIIb-VIIIa | Left | 339 | 2.36 | Cognitive |
| Net 46 | VIIb-VIIIa | Right | 344 | 17.44 | Cognitive |
| Net 47 | VIIb-VIIIa | Right | 354 | 50.56 | Motor |

Network categories were assigned using a 50% spatial-overlap threshold.
